# Deep brain stimulation of the mesencephalic locomotor centre induces stimulation-dependent behavioural states beyond locomotion

**DOI:** 10.64898/2026.09.18.752305

**Authors:** Martina Wilhelm, Johannes Hartig, Teresa Christina Faupel, Dongdong Sun, Saskia Marie Moritz, Jeremy Signoret-Genest, Fereschta Hakimi, Annemarie Sodmann, Felicitas Schlott, Susanne Knorr, Philip Tovote, Chi Wang Ip, Felix Fluri, Michael Klaus Schuhmann, Jens Volkmann, Robert Blum

**Author notes:** These authors contributed equally to this work (Martina Wilhelm, Johannes Hartig). Senior authors (Jens Volkmann, Robert Blum).

## Abstract

Deep brain stimulation (DBS) of the mesencephalic locomotor region (MLR) has been explored to treat gait disturbances. However, clinical outcomes of MLR-DBS have been disappointing overall, with only modest benefits in some individuals and unwanted effects, including anxiety, reported in others. Preclinical studies in several species, by contrast, show that MLR-DBS can improve locomotor and gait deficits, although defensive behaviours in response to stimulation have also been reported in some species. To investigate this discrepancy, we examined frequency- and amplitude-dependent effects of MLR-DBS in healthy rats, combining semi-supervised and unsupervised behavioural analyses. High-frequency, high-amplitude DBS (80–130 Hz), but not low-frequency stimulation (20–60 Hz), elicited hyperlocomotion intermixed with acute defensive-like behaviours. Low-frequency stimulation instead promoted phases of immobility. Consistently, only high-frequency DBS induced region-specific c-Fos expression in the MLR. Complex behaviours, including hyperlocomotion, rearing, tail rattling, and periods of immobility, were most pronounced in animals with the DBS electrode tip localized to the cuneiform nucleus (CnF) of the MLR. Using AAV tracer constructs for bright labelling of CaMKII-positive neurons and their axons, we identified prominent ascending projections from the CnF to the thalamus, substantia nigra pars compacta, zona incerta, hypothalamus, subthalamic nucleus, and central amygdala (CeA). Retrograde tracing confirmed the CnF-to-CeA projection independently. These findings show that the MLR is anatomically connected to higher-order centres linking motor and defensive networks, with DBS frequency and amplitude critically shaping behavioural outcomes that extend beyond locomotion.

## Introduction

Gait disorders are a common feature of neurological conditions affecting the basal ganglia–thalamo-cortical network, such as Parkinson’s disease (PD) and stroke ^1, 2^. This is notable given that locomotion in vertebrates relies on phylogenetically conserved brainstem and spinal circuits ^3, 4, 5^, that are often not directly affected by primary lesions. Indeed, mesencephalic and spinal central pattern generators (CPGs) can generate the alternating rhythmic activity required for stepping, as well as the antigravity postural adjustments needed for walking or running, even after decerebration ^6^. A key structure in this context is the mesencephalic locomotor region (MLR), originally defined functionally as a midbrain site where increasing stimulation intensity elicits a transition from standing to walking and eventually running ^7, 8, 9^. In mammals, the MLR comprises the pedunculopontine nucleus (PPN), cuneiform nucleus (CnF), mesencephalic reticular nucleus (MRN), and their interconnecting fibres ^10, 11^, with functionally and anatomically distinct subregions controlling different aspects of locomotion-related behaviour ^3, 5, 12^. Accordingly, neuromodulation of the MLR with deep brain stimulation (DBS), opto- or chemogenetics, can induce locomotor behaviours and improve gait disturbances in animal models of PD ^13^, incomplete spinal cord injury ^14, 15, 16^, haloperidol-induced catalepsy ^17^, and unilateral stroke ^18, 19, 20^.

Based on these preclinical findings, the PPN has been evaluated as a DBS target for gait disturbances, including freezing of gait (FOG), in PD patients ^21, 22, 23, 24, 25, 26^. This approach rests on the hypothesis that gait disorders reflect a distinct motor circuitopathy arising from abnormal MLR input to lower-order locomotor circuits. Supporting this concept, gait and balance deficits in elderly patients with “higher-level gait disorder” have been linked to lesion or dysfunction of the network connecting motor cortex and the MLR ^27^, and neuronal loss in the MLR is greatest in the PPN and CnF among patients with progressive supranuclear palsy and PD who experience falls ^28^.

However, clinical outcomes of MLR-DBS in the PPN subregion have been inconsistent and overall disappointing, despite modest benefits in some individuals ^21, 29^. This variability has been attributed partly to the precise stimulation site within the MLR, prompting interest in alternative targets such as the CnF ^21, 30^. Converging evidence from animal models and patients with gait disturbances supports the CnF as a promising DBS target ^16, 19, 20, 31, 32, 33^. Recently, it was shown that CnF and PPN stimulation in PD patients with severe gait impairment failed to improve motor, cognitive, or psychiatric outcomes, and instead increased anxiety ^25, 26^. In a separate study, local field potentials recorded from the CnF and PPN were combined with analyses of gait initiation in four patients with FOG. FOG was associated with impaired spatial and temporal precision of neuronal activity, particularly in alpha-band activity patterns within the CnF. CnF stimulation improved gait rhythm, whereas PPN stimulation worsened gait pace ^34^.

Beyond its role in locomotion, MLR activation can also elicit non-locomotor behaviours in animals ^5, 35, 36, 37, 38^. Microinjection of glutamate into the rat CnF, for example, induces freezing, darting, and fast running ^36^, and defensive behaviours following CnF stimulation have been reported consistently in rats ^35, 36, 38, 39^ and pig ^31^. Given the anatomical complexity of the MLR, non-locomotor effects have often been attributed to unintended co-stimulation of neighbouring regions, like the periaqueductal gray (PAG), a key hub for defensive behaviours ^40, 41^. However, complex behaviours after MLR-DBS may instead reflect engagement of higher-order ascending networks ^42, 43^. Indeed, DBS in the MLR can activate higher order regions in the mammalian brain. In rats, ^18^F-FDG-PET confirmed that CnF-DBS activates a broad ascending thalamo-cortical network, before and after stroke ^44^. This is biologically relevant, as high-frequency CnF-DBS triggered anti-inflammatory processes after stroke in the perilesional cortex via modulation of the cholinergic system ^19^. Such findings are in line with neuroanatomical data from humans showing that cuneiform/subcuneiform target regions include thalamic regions, hypothalalmus, basal forebrain, zona incerta and the Globus pallidus, presumably as part of the ascending reticular activating system (ARAS), an essential component of human consciousness ^43^. As locomotion and defensive behaviour are functionally coupled to support state- and context-dependent behavioural states ^45^, it is therefore conceivable that non-locomotor effects of MLR-DBS, often interpreted as off-target effects ^16, 31, 46^, may be mediated by the MLR neurons themselves.

Here, we applied DBS to the MLR in healthy rats and used complementary machine-learning approaches to analyse and segregate voluntary and stimulation-induced behavioural states. We show that MLR stimulation elicits frequency- and amplitude-dependent transitions between defensive and locomotor behaviours, with c-Fos mapping confirming neural activation in both the CnF and PPN. To test whether ascending MLR projections reach brain areas implicated in motor and defensive processing, we developed AAV2-based vectors for anterograde and retrograde tracing with myristoylated GFP in rats. This revealed extensive ascending projections from the CnF to key regions involved in locomotion and defensive behaviour, including the thalamus, zona incerta, hypothalamus, central amygdala (CeA), subthalamic nucleus (STN), substantia nigra pars compacta (SNc), and multiple thalamic nuclei.

## Results

### MLR-DBS reorganizes the multidimensional behavioural landscape of the rat

To characterize behavioural responses following DBS of the MLR, we performed frequency-dependent stimulation in an inverted open-field–like arena. Rats (*n* = 22) were first stimulated unilaterally with high-frequency stimulation (HFS; 130 Hz) at gradually increasing amplitudes to determine each animal’s individual current threshold (Fig. 1a). Behavioural responses during spontaneous activity and under each stimulation condition were recorded with a bottom-view camera for at least 30 seconds. To objectively quantify both DBS-dependent and naturally occurring behaviours, all videos (n = 810) were analysed using keypoint-based tracking in *DeepLabCut* (Mathis et al., 2018) (Fig. 1b).

**Fig. 1.**
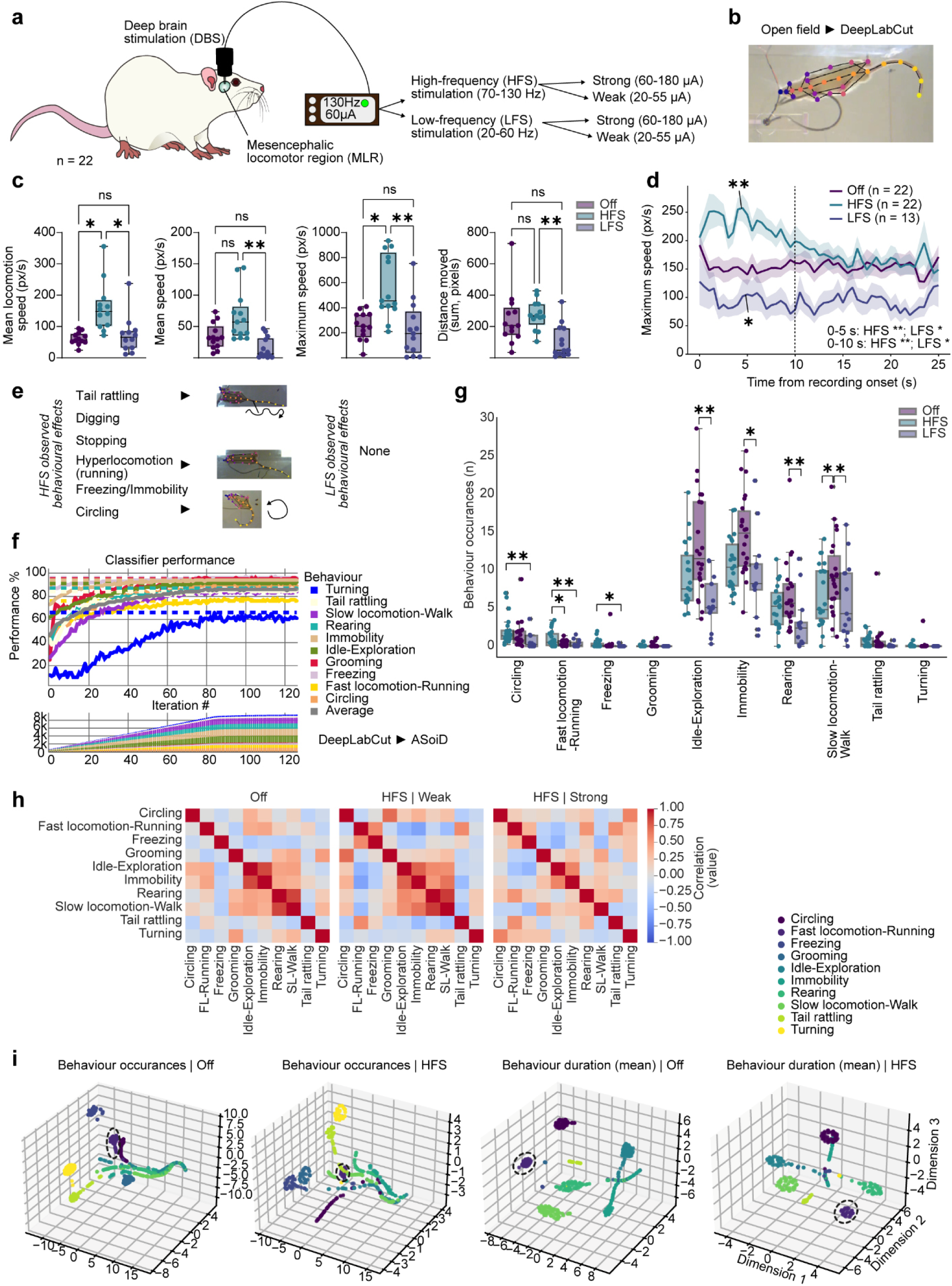
Multidimensional behavioural repertoire induced by electrical DBS of the mesencephalic locomotor region. **a.** Experimental outline. The MLR of Wistar rats (*n* = 22) was stimulated under low and high frequency conditions, with ‘weak’ and ‘high’ stimulation amplitudes as indicated. **b.** Bottom-view in an open arena. The stimulation cable is visible and keypoints from markerless pose estimation of a rat (*DeepLabCut*). **c.** Data from kinematic analysis. Shown are mean locomotion speed (Friedman test, Friedman statistic = 10.17, *p* = 0.0062, two-sided Dunn’s posthocs), mean speed (Friedman test, Friedman statistic = 12.92, *p* = 0.0016, two-sided Dunn’s posthocs), maximum of speed (RM one-way ANOVA, *F* = 10.40, *p* = 0.0007, two-sided Tukey’s posthocs), and distance moved (Friedman test, Friedman statistic = 11.23, *p* = 0.0036, two-sided Dunn’s posthocs), indicating brady-/hypokinesia induced by LFS, enhanced motion induced by HFS. **d.** Immediate onset response shows rapid increase and decrease with HFS (*p* = 0.0022, first 10 s) and LFS (*p* = 0.022, first 10 s), respectively vs. Off (two-sided Wilcoxon signed-rank test, Holm-adjusted *p*-values shown). Data as mean ± s.e.m. **e.** Induced behaviours observed under HFS: tail rattling, digging, stopping, hyperlocomotion (running), freezing or immobility, and circling. LFS: none. **e.** Semi-supervised behavioural classification (*A-SoiD*, ^47^) with and without MLR-DBS displaying classifier performance (F1-score, y-axis). **f.** Number of behaviour occurrences. Classified behaviours (*n* = 810 videos) under control conditions (Off), HFS, or LFS. Categorization included an idle exploration category. The classifier was applied (one Friedman test per behaviour with Nemenyi post-hoc test) and found that we could cross-confirm the running induction (Friedman test, Friedman statistic = 12,04, *p* = 0.002, two-sided Dunn’s posthoc) by HFS (*p* = 0.029) but not LFS (*p* = 0.863). **g.** Correlation matrices show that both weak (20-55 µA) and strong (60-180 µA) HFS leads to similar, yet distinct changes in global behavioural structure and transitions. **h.** t-distributed stochastic neighbour embedding (tSNE; perplexity = 50) shows that for an embedding of both behavioural bout count (left) and mean bout duration (right) in combination with behavioural class and time component, HFS-DBS of the CnF (pooled weak and strong) leads to a profound shift/rearrangement in the behavioural state space of rats. Box plots show median, IQR with whiskers as most extreme data points. Sample sizes: Off: n = 13; HFS: n = 13; LFS: n = 13, if not indicated otherwise.

Kinematic analysis revealed increases in both mean and maximum speed during HFS (70–130 Hz), but not during low-frequency stimulation (LFS; 20–60 Hz; Fig. 1c). Mean speed of locomotion and maximal speed were significantly higher under HFS than under LFS or the Off condition (Fig. 1c). Under LFS, total distance travelled was reduced relative to both HFS and Off. Stimulation onset was followed by an immediate increase in maximal speed within the first 10 seconds under HFS, whereas LFS reduced maximal speed over the same time window (Fig. 1d).

For quantitative analysis of individual behaviours based on the pose-estimation data, we trained a semi-supervised active learning classifier within A-SOiD ^47^. The classifier was trained on sample videos spanning both naturalistic and rater-identified, DBS-induced behaviours, using predefined macro-behavioural categories: turning, tail rattling, slow locomotion (walking), rearing, immobility, idle exploration, grooming, freezing, fast locomotion (running), and circling (Fig. 1e, Supplementary Fig. 1). Supplementary Video 1 illustrates HFS-induced running followed by stop phases that included tail rattling, rearing, and freezing, with the electrode tip histologically confirmed within the CnF subregion of the MLR.

The classifier achieved robust detection accuracy (≥ 75%) for all behaviours except turning (Fig. 1f) and was subsequently applied across all open-arena recordings (*n* = 810 videos). Fast locomotion (running) was increased during HFS but not LFS relative to Off (Fig. 1g), and HFS also showed a stronger tendency toward circling and freezing than LFS. In contrast, LFS was associated with reduced idle exploration (periods without active locomotion or another defined behaviour), immobility, and rearing. Changes in grooming, tail rattling, and turning did not reach statistical significance, although they were observed in individual animals. Mean behaviour duration did not differ significantly across conditions (Off, HFS, LFS), though a trend toward longer immobility bouts under LFS relative to HFS and Off was apparent (Supplementary Fig. 2a).

To examine behaviour duration in more detail, we computed cumulative distribution function (CDF) histograms (Supplementary Fig. 2b–d). Running bouts were longer under HFS (Supplementary Fig. 2b), whereas LFS produced longer walking (slow-locomotion) bouts than either HFS or Off (Supplementary Fig. 2c). Idle exploration duration did not differ across conditions (Supplementary Fig. 2d).

Finally, we asked whether MLR-DBS alters overall behavioural state. We computed correlations between the occurrence of incoming and outgoing behaviours as a measure of how behaviours are linked. LFS did not yield sufficient data points for this analysis, thus we compared Off with HFS only. This revealed distinct, amplitude-dependent correlation patterns for Off versus low- and high-amplitude HFS (Fig. 1h). We then embedded categorized behaviours by frequency and mean duration using t-distributed stochastic neighbour embedding (t-SNE; perplexity = 50) as a qualitative measure of overall behavioural state change. This confirmed that HFS produced a pronounced shift in the multidimensional behavioural repertoire relative to Off, in both the frequency and duration of behaviours (Fig. 1i).

### MLR-DBS-induces interactions between locomotor- and immobile behaviours

To further explore our behavioural data in an unsupervised manner, we trained a *switching linear dynamical system* (SLDS) model to identify behavioural ‘syllables’, discrete motifs of pose dynamics, using *Keypoint-MoSeq* (KPMS) ^48^ (Fig. 2a). The full KPMS model, trained on the entire video dataset (*n* = 810), identified 19 distinct behavioural syllables with a median duration of 350 ms (Supplementary Fig. 3). Individual syllables were labelled based on visual inspection of the trajectory videos. Relative to *idle exploration* (syllable 0), syllable 1 was classified as ‘*exploration / brief forward locomotion*’, syllable 2 and 6 as two distinct immobile states, syllable 9 as ‘*turn left and go’*, syllable 10 as ‘*naturalistic locomotion*’, syllable 11 as ‘*circle and go*’, and syllable 15 as ‘*stimulation-induced hyperlocomotion*’, which also included jumping.

**Fig. 2.**
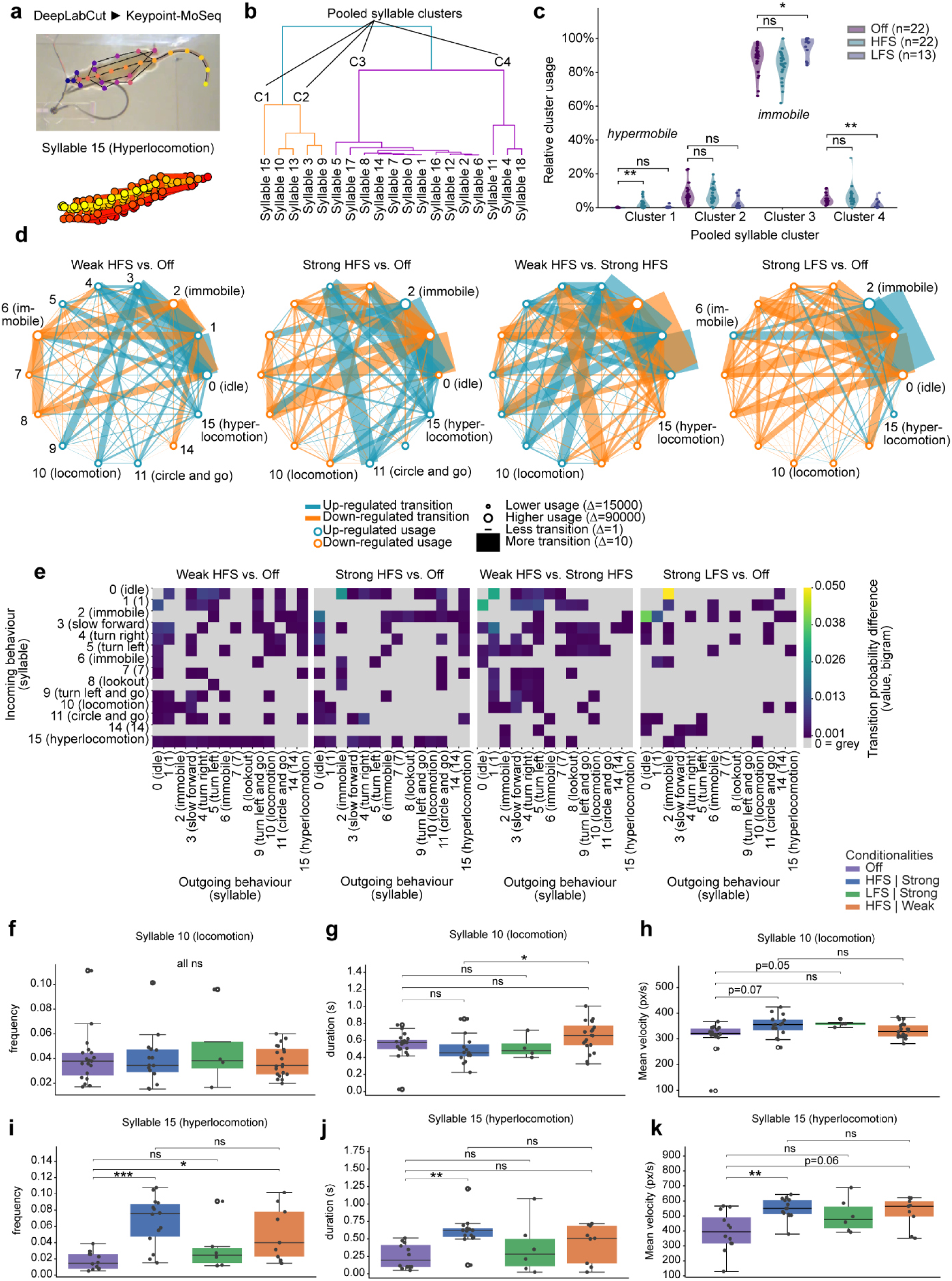
Low- and high-amplitude HFS-DBS of the MLR differentially affects naturalistic and induced locomotor behaviours. **a.** Experimental outline. Behavioural videos (*n* = 810 videos, *n* = 22 rats) were pose-tracked using *DeepLabCut* and analysed with *Keypoint-MoSeq* for unsupervised identification of behavioural syllables (example: syllable 15, hyperlocomotion). See Supplementary Fig. 3 for full set. **b.** Syllable dendrogram depicting differences between identified syllables based on median trajectories. **c.** Relative cluster usage shows increased hypermobile cluster usage with HFS (Friedman test per cluster, cluster 1: Friedman statistic: 16.21, *p* = 0.00030) and immobile cluster usage with LFS (cluster 3: Friedman statistic: 17.08, *p* = 0.00020) vs. Off. **d.** Node and network plots depicting the behaviour usage difference (node size) and transition difference (line thickness) between the syllables for the different conditionality comparisons. Up-(turquoise) and downregulated (orange) usage and transitions between behaviours and difference categorization values are given. **e.** Matrix plots showing the transition probability difference of existing behaviours (incoming behavior) and the subsequent behaviour (outgoing) for indicated comparisons. Spectral procrustes analysis of difference in global structure of transitions were all non-significant (Bonferroni corr. *p*-values all = 0.99). **f-h.** Syllable 10 (naturalistic walking, locomotion) analysis. Frequency does not change (Kruskal-Wallis test: H-statistic: 0.032, *p* = 0.9988), while duration only increases with low-amplitude vs. high-amplitude HFS (*p* = 0.032), but not Off (Kruskal-Wallis test: H-statistic: 7.63, *p* = 0.0482). Mean velocity (Kruskal-Wallis test: H-statistic: 10.04, *p* = 0.0436) shows trends for increase with both high-amplitude HFS (*p* = 0.0664) and LFS (*p* = 0.0515). **i-k.** Syllable 15 (hyperlocomotion) analysis. Frequency increases with both high-(*p* = 0.0006) and low-amplitude HFS (*p* = 0.0369), but not with high-amplitude LFS (*p* = 0.9926; Kruskal-Wallis test: H-statistic: 15.46, *p* = 0.0001). Duration only increases with high-amplitude HFS (*p* = 0.0013; Kruskal-Wallis test: H-statistic: 13.70, *p* = 0.0008). Mean velocity increases with high-amplitude HFS (*p* = 0.0064) and trend for low-ampl. HFS (*p* = 0.0562). Syllable statistics all as Kruskal-Wallis tests with animal-level clustered permutations and post-hoc Dunn‘s z-test (Holm-adj. sign. depicted) per syllable across groups. Box plots show median, IQR with whiskers as most extreme data points. Sample sizes: Off: *n* = 22; HFS: *n* = 22; LFS: *n* = 13, if not indicated otherwise.

To characterize how MLR-DBS reorganizes the structure of naturalistic and induced behaviours, we constructed a syllable dendrogram based on Euclidean distance between syllable trajectories (Fig. 2b), yielding four clusters: cluster 1 comprised syllable 15 alone (“hyperlocomotion”); cluster 2 comprised predominantly active behaviours; cluster 3 comprised various low-mobility behaviours; and cluster 4 comprised active turning behaviours. We then computed relative cluster usage under LFS, HFS, and Off (Fig. 2c). Consistent with the findings above, the hypermobile cluster was enhanced by HFS relative to Off, while the immobile cluster was enhanced and active turning behaviours were reduced by LFS.

To visualize transition frequencies between syllable pairs, we generated node and network plots (Fig. 2d; Supplementary Fig. 4a for condition-specific plots). Under low-amplitude (“weak”) HFS, diverse syllables (mostly in cluster 1, 3, 4) showed upregulated transitions into hyperlocomotion (syllable 15) and naturalistic locomotion (syllable 10). Transitions involving immobility-related syllables (2, 6, 7, 8, 14, mostly cluster 3) were downregulated (Fig. 2d, leftmost). High-amplitude (“strong”) HFS additionally promoted direct transitions from idle exploration into an immobile state (syllable 2), from a diverse set of syllables including locomotion and hyperlocomotion (cluster 1,2 with locomotion and hyperlocomotion) and transitions within cluster 4 (syllable 4 and 11 “circle and go”) (Fig. 2d, second from left). Direct comparison of low-versus high-amplitude HFS showed reduced transitions into both immobility (syllables 2 and 6) and hyperlocomotion (syllable 15) at high amplitude, indicating a dose-dependent relationship (Fig. 2d, second from right). High-amplitude LFS, by contrast, strongly enhanced transitions between idle exploration and immobility syllables 1, 2 and downregulation of virtually all syllables in the clusters 1,2,4 (Fig. 2d, rightmost).

We next generated transition matrices as a complementary measure of behavioural transition probabilities (Fig. 2e; Supplementary Fig. 4b for native transition matrices). A spectral Procrustes analysis comparing the eigenvalues of these matrices revealed no significant differences between conditions. However, examining specific transition-probability changes, high-amplitude LFS produced a bradykinetic effect, increasing the probability (∼5%) of transitioning from idle exploration into immobility (Fig. 2e, rightmost panel).

Finally, analysis of in-syllable kinematics (Fig. 2f–k) showed that weak HFS, relative to strong HFS, preferentially promoted naturalistic locomotion (Fig. 2g), while high amplitude increased mean velocity under both LFS and HFS (Fig. 2h). As expected, HFS increased the frequency (Fig. 2i), total duration (Fig. 2j), and mean velocity (Fig. 2k) of hyperlocomotion; the frequency and mean velocity of hyperlocomotion were likewise increased under weak HFS (Fig. 2i,k). The kinematics of syllable 11 (“circle and go”) were also significantly regulated, with strong HFS increasing its frequency and velocity, and weak HFS increasing its duration (Supplementary Fig. 4). High-amplitude HFS increased the frequency of hyperlocomotion/running specifically, without affecting the frequency of naturalistic locomotion (Fig. 2g). Effects on duration were dissociable: increased hyperlocomotion duration was detectable only with high-amplitude HFS (Fig. 2h), whereas naturalistic locomotion duration was affected only by low-amplitude HFS (Fig. 2i).

Taken together, unsupervised analysis of behaviour showed that high-frequency MLR stimulation in rats has amplitude-dependent effects on naturalistic locomotion (walking) and acutely promotes hyperlocomotion (running) for several seconds, whereas high-amplitude, low-frequency stimulation promotes transitions into immobility-like states.

### MLR-DBS elicits region- and frequency-specific c-Fos

To further characterize the local biological effects of MLR-DBS, *n* = 17 Wistar rats were stimulated in their home cages with either high-frequency stimulation (HFS, 130 Hz) or low-frequency stimulation (LFS, 40 Hz). As a biological marker, we assessed *de novo* synthesis of c-Fos protein (Fig. 3a), an activity-dependent transcription factor whose expression is upregulated in response to DBS ^20, 49^. For controls, the stimulator and electrode were implanted and the cable connected, but the stimulator was not switched on. Following our previous work ^50^, experiments were conducted in the home cage to minimize context-dependent c-Fos activation in freely behaving animals. All animals received stimulation for 60 minutes, and brain tissue was collected 90 minutes after stimulation onset. c-Fos-labelled images were quantified using a deep learning model ensemble on tile-imaged whole-brain slices with *deepflash2* ^50, 51^.

**Fig. 3.**
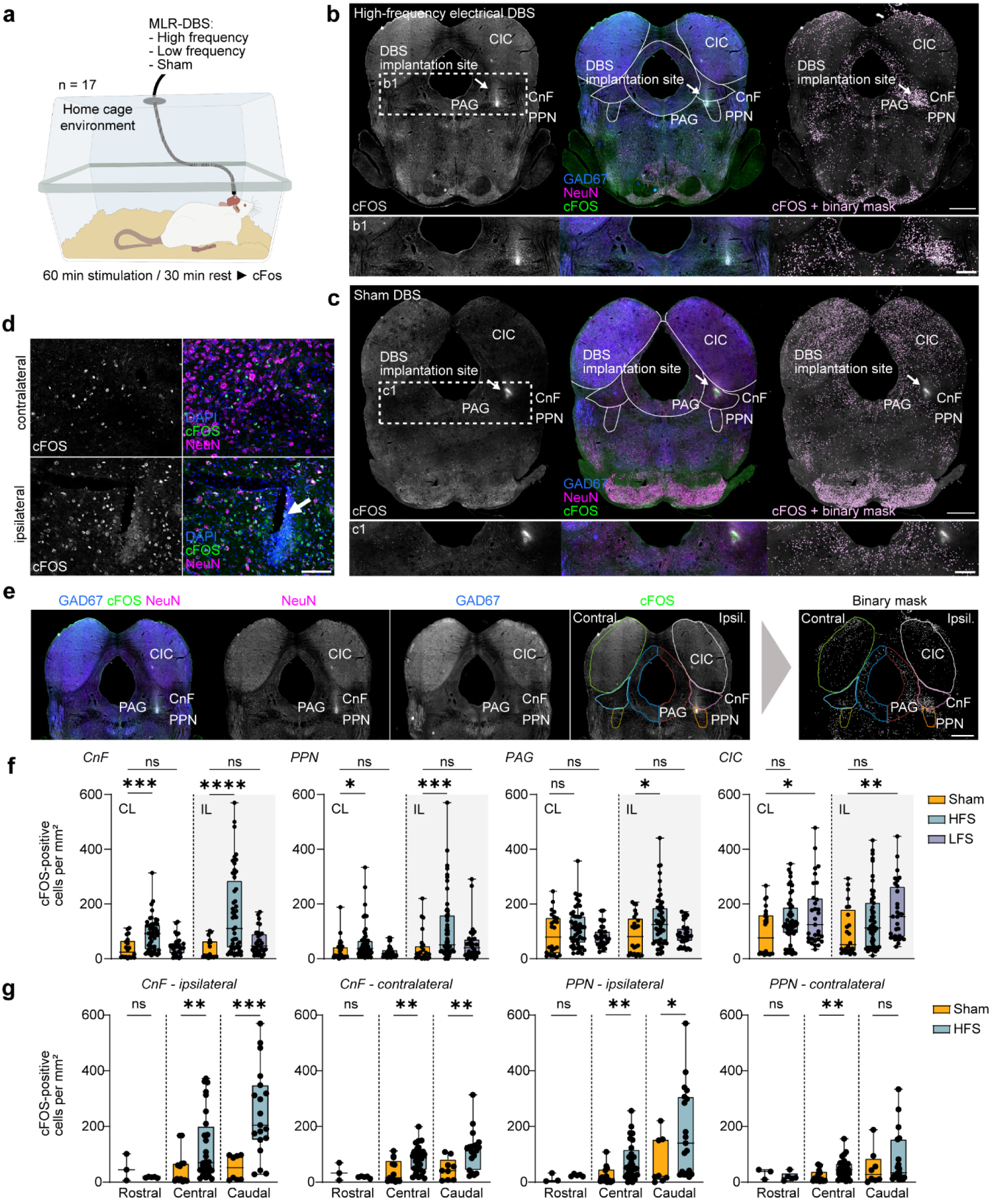
High-frequency deep brain stimulation of the MLR induces region- and hemisphere-specific neuronal activation. **a.** Experimental schematic: Wistar rats (n = 17) with MLR-DBS were stimulated under high-frequency (HFS), low-frequency (LFS) or sham condition for 60 min followed by 30 min rest before c-Fos quantification. **b.** Representative coronal sections after high-frequency DBS showing c-Fos expression (left), merged immunostaining for GAD67, NeuN and c-Fos (middle), and binarized c-Fos signal (right). Scale, 1 mm. Insets (b1) show higher magnification of the stimulated region. Scale, 500 µm. **c.** Representative coronal sections after sham stimulation showing c-Fos expression (left), merged immunostaining (middle) and binarized c-Fos signal (right). Scale 1 mm. Insets (c1) show higher magnification of the corresponding region. Scale, 500 µm. Arrows in b-c) indicate electrode position. **d.** Higher magnification images comparing contralateral and ipsilateral hemispheres, showing c-Fos signal (left) and merged staining (DAPI, c-Fos, NeuN; right). Arrow indicates electrode position. Scale, 100 µm. **e.** Region-of-interest definition and signal extraction workflow showing individual channels (GAD67, NeuN, c-Fos) and anatomical segmentation of CnF, pedunculopontine nucleus (PPN), periaqueductal gray (PAG) and inferior colliculus (CIC), followed by binary mask generation. Scale, 1 mm. **f.** Quantification of c-Fos-positive cells per mm² across regions (CnF, PPN, PAG, CIC) and hemispheres (contralateral, CL; ipsilateral, IL) under sham (orange), HFS (blue) and LFS (purple) conditions. Statistical significance was assessed using the Kruskal Wallis test followed by Dunn’s multiple comparison. **g.** Rostrocaudal analysis of c-Fos-positive cell density in CnF and PPN for contralateral and ipsilateral hemispheres under sham (orange) and HFS (blue) conditions. Statistical significance was assessed using unpaired t-tests with Welch’s correction or Mann–Whitney U tests, as appropriate. In e-f.: boxplots show median, IQR, and whiskers indicate minimum–maximum values. Individual data points represent single brain slices (5-8 slices per animal). Number of slices analysed: sham, n = 24; LFS, n = 31; HFS, n = 52. Animal sample sizes were: sham, n = 4; LFS, n = 4; HFS, n = 9.

Following HFS, a marked increase in c-Fos-positive cells was observed in the ipsilateral MLR by tile microscopy (Fig. 3b) and confirmed at higher resolution by confocal microscopy (Fig. 3c). To quantify the density of c-Fos-positive neurons (cells/mm²) and their distribution across defined anatomical regions, deep learning– based segmentations of c-Fos labels, were analysed in the MLR subnuclei CnF and PPN, as well as in adjacent medial and dorsal regions, the periaqueductal grey (PAG) and central nucleus of the inferior colliculus (CIC), in both hemispheres (Fig. 3d).

This quantitative analysis revealed a significant increase in the density of c-Fos-positive neurons in both the ipsilateral (stimulated) and contralateral CnF and PPN following HFS compared to sham stimulation, with a more pronounced effect on the stimulated side (Fig. 3e). The ipsilateral PAG also showed a significant increase in c-Fos-positive neurons after HFS. In the CIC, by contrast, both hemispheres showed a significant increase in c-Fos-positive cells following LFS compared to sham stimulation.

We next examined the rostro-caudal distribution of c-Fos-positive neurons in the CnF and PPN, analysing rostral, central, and caudal subdivisions after sham stimulation and HFS. In the CnF, both the central and caudal subdivisions showed a significant increase in c-Fos-positive neurons after HFS compared to sham, on both the ipsilateral and contralateral sides (Fig. 3f).

Together, these findings indicate that c-Fos activation following local MLR-DBS spreads along the rostro-caudal axis within MLR subregions, engages the contralateral MLR, and extends into the PAG.

Behavioural responses following HFS of the MLR and its subnuclei in the home cage were heterogeneous, with individual animals frequently exhibiting multiple behavioural categories (Supplementary Fig. 5). Across all MLR stimulations, immediate locomotion was the predominant response (10/11 animals), followed by rearing (6/11). Intermittent stop-and-go behaviour and clockwise rotation were each observed in 4/11 animals, and jumping in 3/11. Less frequent behaviours included digging, freezing, and stopping with tail rattling (2/11) (Supplementary Fig. 5).

In individual animals, electrode placement appeared to shape the behavioural response. For example, in an animal with the electrode tip in the CnF, high-frequency DBS induced immediate locomotion, followed by cessation of movement, tail rattling, and digging, and then, within approximately 30 seconds, freezing (Supplementary Fig. 6, compare with supplementary video 1 in the open arena). In this animal, c-Fos labelling was more prominent ipsilaterally, particularly in the CnF, PPN, and PAG, but not in the adjacent inferior colliculus (Supplementary Fig. 6b, c), with c-Fos expression preferentially increased within the CnF both rostral and caudal to the stimulation site. In another animal, stimulation within the electrode tip in the PPN (Supplementary Fig. 7) also elicited immediate locomotion, but this was followed by clockwise rotation, rearing, and stop-and-go behaviour. In this case, c-Fos expression was elevated primarily near the electrode tip, with minimal activation in surrounding regions (Supplementary Fig. 7b, c). Together, these observations indicate that both the behavioural response to DBS and the pattern of neuronal activation varied with electrode position. Expert-rated behaviours and electrode positions for individual animals, mapped onto the corresponding brain areas, are shown in Supplementary Fig. 8.

### Ascending fibres from cuneiform nucleus neurons of the rat target multiple higher-order brain regions

We previously showed using [18F]-FDG-PET that high-frequency CnF-DBS activates a widely distributed thalamo-cortical network, before and after stroke ^44^. As such network-level metabolic changes typically serve as a proxy for regional neuronal and synaptic activity, we next investigated the projection targets of ascending axonal fibres from the CnF directly. Because most viral tracing vectors do not permit bright axon labelling in the rat, we generated neuron-specific fibre-tracing constructs based on AAV2, the *CaMKII(1.3)* promoter ^52^ and a myristoylated GFP (myrGFP). To generate myrGFP, we fused to an N-terminal myristoylation/palmitoylation sequence from LCK, via a flexible GGSGG motif, to GFP ^53, 54^ resulting in *AAV2-CamKII(1.3)-myrGFP* (Fig. 4a). This modification is particularly suited to label axon fibres and presynapses and was therefore used for anterograde tracing ^54, 55, 56^. A second construct (*AAV2-CamKII(1.3)-myrGFP-P2A-NLS-tdTomato*) co-expressed myrGFP with nuclear tdTomato (Fig. 5a). This construct was less bright but allowed identification of the cell bodies expressing myrGFP and was therefore used for retrograde tracing.

**Fig. 4.**
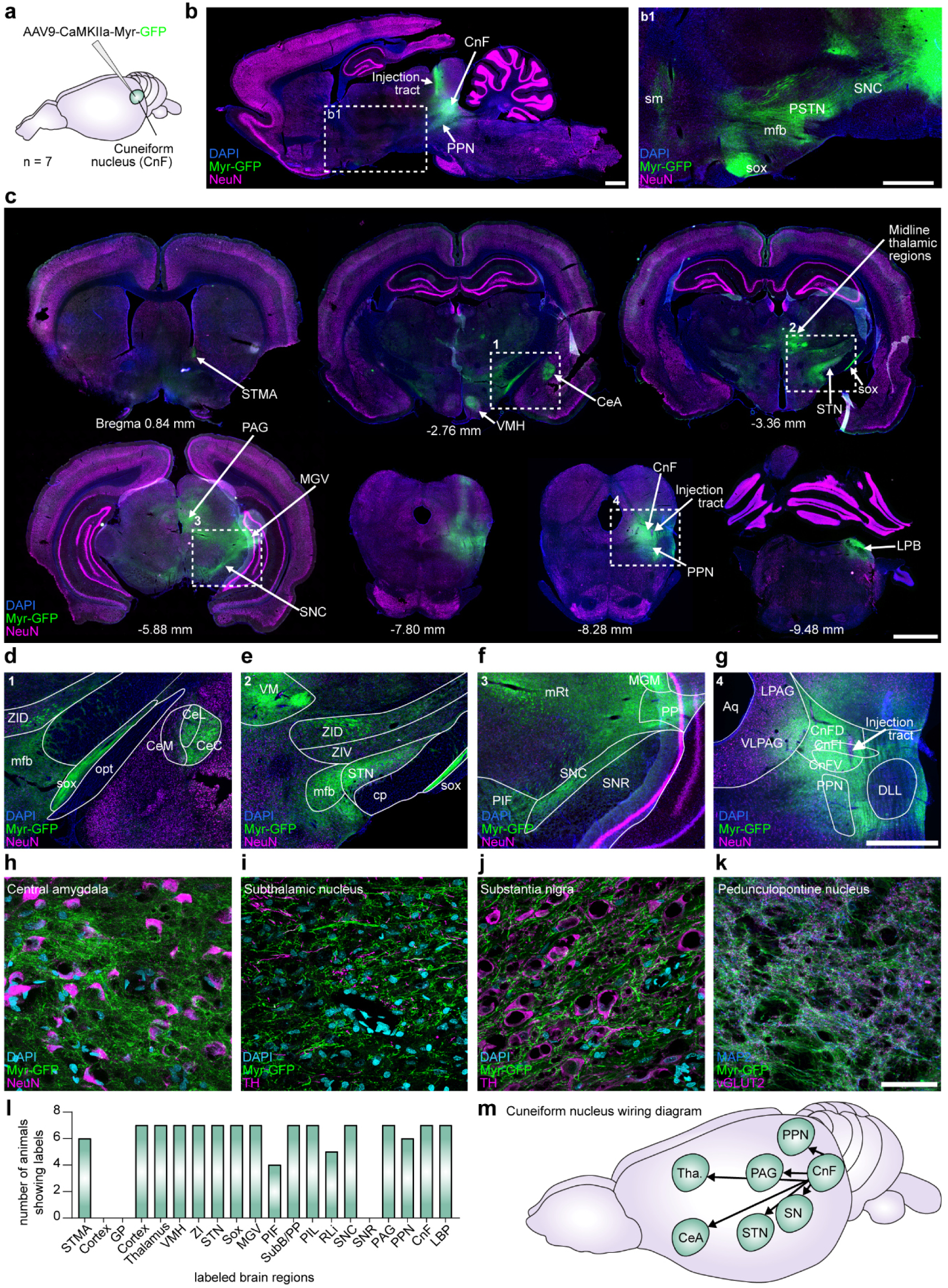
Anterograde connectivity of the cuneiform nucleus in rats with motor and non-motor areas. **a.** Schematic of viral tracing strategy: AAV-CamKII-Myr-GFP was injected into the CnF to label glutamatergic projections (*n* = 7 Wistar rats). **b.** Representative sagittal overview showing injection site and widespread projections from the CnF across the brain. Scales, 1 mm. **c.** Coronal sections illustrating dense anterograde projections of CnF neurons to multiple target regions, including basal ganglia, thalamic and brainstem areas. Scale, 2 mm. **d-g.** Higher magnification views of projection targets, including CeA, ZI, different thalamic nuclei and other subcortical regions. Scale, 1 mm. **h-k.** Confocal high-resolution images showing fibre signals in selected regions. **l.** Number of animals showing fibre labels per atlas brain region of interest. **m.** Summary scheme illustrating major projection targets of labelled CnF neurons. CeA, central amygdala; CnF, cuneiform nucleus; PAG, periaqueductal grey; PPN, pedunculopontine nucleus; SN, subthalamic nucleus; STN, substantia nigra pars compacta, Tha., thalamus, ZI, zona incerta. Scale, 50 µm.

**Fig. 5.**
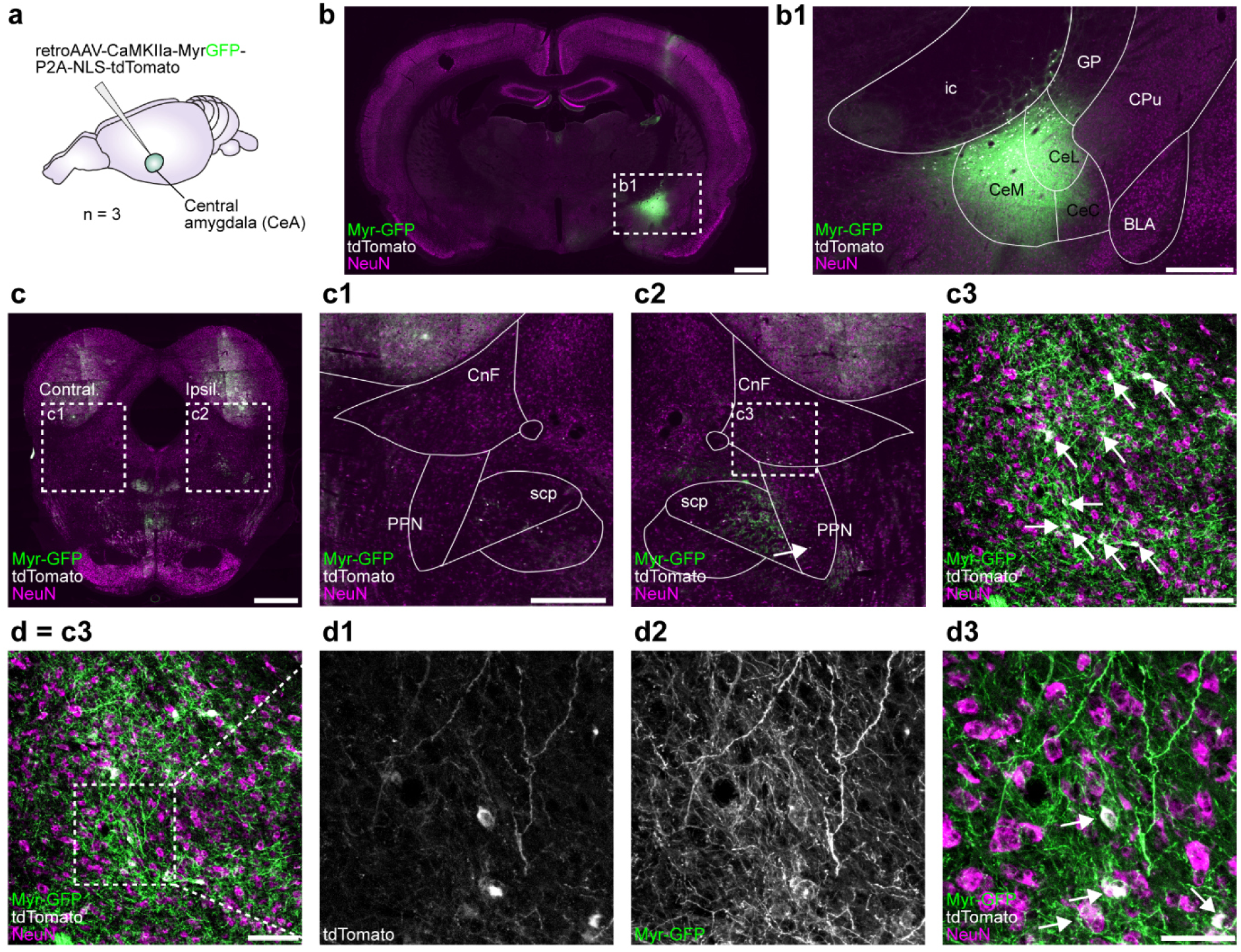
Reciprocal connectivity of the cuneiform nucleus with the central amygdala in rats. **a.** Retrograde viral tracing strategy: retroAAV-*CamKII(1.3)*-MyrGFP, P2A-linked to a nuclear tdTomato was injected into the central amygdala (CeA) (*n* = 3 Wistar rats). **b.** Representative overview showing injection site in CeA. MyrGFP (green), tdTomato (white), and NeuN (magenta) labels are shown. Scales, b = 1 mm, b1 = 500 µm. **c.** Coronal sections illustrating retrogradely labelled neurons connecting the MLR and the CeA. Scales, c = 1 mm, c1/c2 = 500 µm, c3 = 100 µm. **d.** Confocal high-resolution images of tdTomato-labelled neurons and associated fibers in the CnF, demonstrating reciprocal connectivity between both regions. Scales, d/c3 = 100 µm, d1-d3 = 50 µm.

Wistar rats (*n* = 7) were unilaterally injected at stereotactic coordinates targeting the centre of the CnF, a site at which DBS mediates locomotor responses (this study and ^15, 16, 20^) and to activate higher-order brain centres via MLR-DBS ^44^. MyrGFP expression was robust and bright, allowing direct visualisation of fibres without additional immunostaining (Fig. 4b). In sagittal sections, the injection track was clearly identifiable, showing myrGFP fluorescence along its trajectory. MyrGFP labelling further revealed an extensive ascending fibre network originating from the MLR. Fibres could be traced continuously from the MLR through the substantia nigra pars compacta (SNc) and parasubthalamic nucleus (PSTN), following the medial forebrain bundle (mfb). From there, projections extended ventrally toward the supraoptic decussation (sox) and rostrally, where signal intensity faded over a short distance before reappearing more distinctly within the stria medullaris of the thalamus (sm).

We collected 40 µm coronal cryosections and screened for fibre signal using tile microscopy (Fig. 4c). Descending fibres were observed in the ipsilateral lateral parabrachial nucleus (LPB). Ascending projections extended predominantly ipsilaterally to the SNc, the medial geniculate nucleus (MGV), the periaqueductal grey (PAG), the subthalamic nucleus (STN), the zona incerta (ZI), the sox, thalamic regions, the central amygdala (CeA), the ventromedial hypothalamic nucleus (VMH), and, most rostrally, the bed nucleus of the stria terminalis (STMA, Fig. 4c). Several regions, particularly the SNc, sox, and thalamic regions, also showed a very weak contralateral signal (Fig. 4c).

Within the CeA, labelling was restricted to the lateral and capsular divisions (CeL, CeC), excluding the medial division (CeM). The sox showed dense fibre labelling (Fig. 4d). Medial to the STN, the medial forebrain bundle (mfb) showed a bright projection pattern (Fig. 4e). Dorsal to the STN, the dorsal and ventral zona incerta (ZID, ZIV) were labelled. Dense labelling was also observed in the ventromedial thalamic nucleus (VM) (Fig. 4e). The signal in the SNc extended dorsolaterally toward the peripeduncular nucleus (PP) and medial geniculate nucleus (MGM), and ventromedially toward the parainterfascicular nucleus of the ventral tegmental area (PIF) (Fig. 4f). From the MGM, fluorescence extended medially into the mesencephalic reticular formation (mRt). Signal passing through the SNc largely avoided the substantia nigra pars reticulata (SNr), with only sparse labelling in the dorsolateral region (Fig. 4f). In the MLR, a strong signal was concentrated across all three subdivisions of the cuneiform nucleus (CnFD, dorsal; CnFI, intermediate; CnFV, ventral), from where labelling spread medially into the adjacent lateral and ventrolateral periaqueductal grey (LPAG, VLPAG) (Fig. 4g). Ventrally, a gradually weakening signal extended into the PPN, entirely sparing the dorsal nucleus of the lateral lemniscus (DLL) entirely (Fig. 4g).

We used high resolution confocal microscopy to document the fibers in specific target regions. MyrGFP-positive fibres were dense in the central amygdala (Fig. 4h, the STN (Fig. 4i) and SNC (Fig. 4j). In the STN, these fibres were observed alongside tyrosine hydroxylase-positive (TH) fibres, originating from the SNc (Fig. 4i), whereas in the SNc, MyrGFP-positive fibres were interspersed with TH-positive neurons (Fig. 4j).

Among the regions analysed, the CeA, STN, and SNc showed consistent myrGFP-positive labelling across all animals, alongside several additional regions (Fig. 4l).

The ascending projection pattern was partially consistent with the literature from tracing in the rat and cat and tractography in human post-mortem brains ^35, 42, 43, 57^. Notably, however, anterograde tracing identified the central amygdala as a previously rarely reported CnF target region ^35^. To confirm this connection, we performed retrograde tracing (*n* = 3 Wistar rats) using rAAV2 encoding myrGFP and nuclear tdTomato (Fig. 5a). This vector was injected unilaterally into the CeA of three Wistar rats (5 × 10⁹ genome copies (gc) in 35 nl) (Fig. 5a). MyrGFP was robustly and brightly expressed in the CeA (Fig. 5b), with signal spanning the lateral, medial, and capsular divisions (CeL, CeM, CeC). tdTomato-positive neurons were likewise observed in all three divisions. Both myrGFP and tdTomato sparsely labelled the adjacent border of the globus pallidus (GP), while no signal was observed in the basolateral amygdala (BLA) or caudate putamen (CPu). In the MLR, the ipsilateral side showed tdTomato-positive neurons not only in the superior cerebellar peduncle (scp) but also in the CnF and the ventral PPN (Fig. 5c). High-resolution confocal imaging revealed a bright myrGFP-positive fibre network interspersed with tdTomato-positive, NeuN-co-labelled neurons in the CnF (Fig. 5d). On the contralateral side, tdTomato-positive labelling was restricted to the scp.

## Discussion

DBS of the mesencephalic locomotor region (MLR) has been explored as a treatment for gait disturbances in Parkinson’s disease, stroke, and spinal cord injury ^16, 21, 22, 24, 25, 26, 29, 46^. This approach rests on the rationale that MLR neurons couple descending locomotor pathways with ascending projections to coordinate spinal circuits and higher-order motor control ^4, 5, 6^. Clinical efficacy, however, remains variable, and stimulation-induced side effects such as oscillopsia, sensory phenomena, and increased anxiety suggest that MLR stimulation engages behavioural systems beyond locomotion ^4, 5, 6, 24, 25, 26, 29, 46, 58^.

Animal studies, often in rats, support this view. MLR stimulation robustly drives locomotion ^14, 15, 17, 20, 59, 60^, but can also elicit freezing, escape-like behaviour, and heightened arousal depending on species, stimulation parameters, and target ^10, 31, 35, 36, 37, 38, 39^. Cell-type-specific MLR lesions have even been linked to cataplexy and episodic immobility of gait ^10^. The MLR therefore appears to act as a multifunctional hub integrating motor and defensive programs, and may contribute to higher-order functions attributed to the ascending reticular activating system (ARAS), at least in humans ^43^.

Where and how DBS must be applied to promote voluntary gait remains unresolved. Most animal studies targeting the MLR have investigated acute, stimulation-induced locomotion. In rats, for instance, a functional hotspot inducing immediate locomotion under disease conditions has been precisely localized to a volume around the CnF and PPN (stereotaxic coordinates AP: ∼−8.2 to −7.6, DV: ∼−4.8 to −5.7, MV: −2 to −2.7, (according to ^61^) ^14, 15, 16^, where 50 Hz stimulation delivered via a ∼250 µm electrode reliably induced locomotor responses ^14, 15, 16^. In previous work from our group, the same CnF hotspot was targeted before and after stroke using a range of frequencies (30–130 Hz) and amplitudes ^33^. Increased gait velocity was achieved with either low frequency and high amplitude (e.g., 30 Hz, 120 µA) or high frequency and low amplitude (e.g., 130 Hz, 60 µA) stimulation, with low-amplitude, high-frequency stimulation proving particularly effective for recovering gait after unilateral stroke ^19, 20, 44^. In humans, by contrast, low frequencies (∼10–40 Hz) are typically used clinically, while higher frequencies (60–130 Hz) have been reported as ineffective or even gait-worsening ^21, 23, 25^.

In this study, c-Fos expression, used as a proxy for recent neuronal and network activation, was induced by MLR-DBS preferentially in the CnF and PPN, predominantly ipsilaterally and independent of exact electrode position. Given the tight CnF–PPN interconnection ^60^ and our tracing data showing a fibre-rich projection field of the CnF neurons within the MLR itself, a small stimulating electrode is likely to engage both structures simultaneously. In one animal with an electrode precisely placed in the CnF hotspot, home-cage monitoring captured a clear behavioural sequence of running, tail rattling, digging, and freezing, with c-Fos engaged across the CnF, PrCnF, and PPN, consistent with dynamic switching between behavioural states rather than a single, fixed motor output.

Off-target spread to neighbouring structures, particularly the periaqueductal gray, is a common explanation for defensive-like behaviours reported for MLR-DBS ^31, 39^. However, as discussed above, CnF neurons give rise to many other higher order brain regions, including the central amygdala or the zona incerta, which also integrate emotional, sensory, and autonomic signals to coordinate defensive and affective behaviour. Thus, MLR stimulation may not simply drive motor output, but simultaneously broadcast state signals to circuits governing fear, arousal, and motivational valence. High-frequency stimulation preferentially induced active states (locomotion, hyperlocomotion, defensive-like responses), while lower frequencies promoted immobility. These frequency-dependent transitions, together with congruent c-Fos patterns, support a model in which MLR circuits are part of state-switching networks.

### Limitations of the study

Sparse viral labelling revealed a dense MLR-internal projection pattern but could not resolve whether individual CnF neurons within the locomotor hotspot divergently target both defensive and motor networks via axon collaterals, a question that will require single-neuron tracing. The *CaMKII(1.3)* promoter is preferentially active in glutamatergic neurons ^52, 62^ and the targeted neurons were NeuN-positive and GAD67-negative. We therefore infer that our tracing virus predominantly targeted glutamatergic projection neurons, although we lack direct confirmation of this. Confocal microscopy identified synapse-like varicosities in target regions (e.g., CeA, SNc), but whether these synapses are functionally engaged by DBS remains unknown.

Electrode position varied slightly despite consistent targeting. For 3R reasons, animals with post-mortem evidence of electrode mal-positioning were not excluded from this study, as prior MLR-DBS studies have also reported locomotion induction outside the currently defined anatomical boundaries of the CnF/PPN ^5, 8^. Precisely mapping electrode positions across studies onto the Paxinos and Watson atlas also remains difficult, limiting cross-study comparison. A further critical translational question is which structures MLR-DBS would engage once CnF/PPN neurons are lost to neurodegeneration ^28^, a question difficult to address in current PD animal models ^63^.

Together, our findings position the MLR as an integrative node linking motor and emotional systems via divergent ascending projections, offering a possible explanation for the inconsistent clinical and experimental outcomes reported for MLR-DBS. Variability between studies may reflect the inherent functional architecture of MLR circuits rather than technical limitations or suboptimal electrode targeting. Improving clinical efficacy will likely require moving beyond anatomical targeting alone, toward circuit- and state-specific neuromodulation strategies ^64, 65^. This includes more speculative possibilities, such as leveraging plasticity within MLR–forebrain loops to bias network dynamics toward locomotor-dominant states while minimizing recruitment of defensive circuitry.

## Methods

### Animals

All experiments were conducted in accordance with European Union guidelines and were approved by our Institutional Animal Care and Use Committee and by the local authorities (Regierung von Unterfranken; approval numbers RUF55.2.2-2532-2-1309, −2-1800, and −2-1602). Experiments were performed in accordance with the ARRIVE guidelines.

Male Wistar rats (176–200 g; Crl:WI; #003WISTAR, Charles River Laboratories, Sulzfeld, Germany) were housed in groups of three (type IV cage) under a 14/10 h light–dark cycle (10–45 lux), at an ambient temperature of 20–24°C and humidity of 45–65%. Environmental enrichment, consisting of paper pulp and egg boxes, was provided twice weekly. Food and water were available ad libitum. Animals were allowed to acclimatise for at least 12 days prior to surgery. Surgeries were performed once animals reached a body weight of ∼ 270–310 g, matching the weight range used to construct the rat brain stereotaxic atlas ^61, 66^. After electrode implantation, animals were housed individually (type III cage). Rats were pathogen-free, and animals were randomly assigned to experimental and control groups.

### Viral vector cloning and virus production

For cloning of tracing vectors, pAAV-CaMKIIa-WPRE-BGHpA was used as backbone (#51087, Addgene). The vector carries the CamKII(1.3) promoter ^52^. Expression cassettes were designed *in silico* and synthesised by ATG:biosynthetics (Merzhausen, Germany). To clone the vector pAAV2-CamKII(1.3) MyrGFP, an insert encoding Myr-GFP was cloned into the backbone via BamHI and EcoRI. The insert consisted of an artificial Kozak sequence (CGCCACCATG), followed by the LCK myristoylation/palmitoylation consensus sequence (MGCVCSSNPEDD), a GGSGG-linker, and eGFP (ΔATG) ^54^. A NheI restriction site was introduced before the stop codon. To clone pAAV2-CamKII(1.3) MyrGFP-P2A-NLS-tdTomato, a synthesised P2A-NLS-tdTomato insert was introduced via NheI and EcoRI.

Following cloning and sequence verification, AAVs were produced by the in-house viral production facility at the Institute of Clinical Neurobiology, University Hospital Würzburg. Viral particles were generated using the helper plasmids pHGT1-Adeno1, AAV2retro\cap, or pAAV2/9n to obtain AAV2/9 or retroAAV2 pseudotypes. Viral titres (genome copies / ml) were quantified by qPCR using external standard curves and primers targeting the WPRE or WPRE-BGH sequence ^67^. Viral vectors were diluted in 1× PBS. Virus preparations were stored in PBS at −80°C in 4 µl aliquots. During surgery, viral vector aliquots were kept on ice.

### Stereotaxic surgeries

Animals (mean body weight 294 g) were weighed prior to surgery. Preoperative analgesia was administered as Tramadol (30 mg/kg body weight [BW], s.c.) or Metamizole (200 mg/kg BW, s.c.). Anaesthesia was induced with 4% isoflurane (Iso-Vet) in 0.8 L/min oxygen in an induction chamber and maintained at 3% after the animal was placed in a stereotaxic frame (Stereotaxic Alignment System 1900, Kopf Instruments). Body temperature was maintained at 37°C using a rodent warming system with a heating pad and rectal thermal probe (Stoelting). Eye ointment (Bepanthen, Bayer) was applied to prevent corneal drying during surgery, and the skin was disinfected with Octenisept (Schülke). Lidocaine hydrochloride 2% (Bela-Pharm GmbH, 0.2 ml) was injected subcutaneously as a local anaesthetic along the planned incision. After a toe-pinch reflex test confirmed adequate anaesthesia, the skull was exposed and isoflurane was reduced to 1.8%.

The bregma–lambda difference was measured using the Centering Scope 40× (Kopf). The Stereotaxic Alignment Indicator tool (Kopf) was adjusted to the measured difference and positioned on bregma and lambda to level the sagittal plane. To level the coronal plane, the indicator tool was adjusted to 6 mm and rotated by 90° to measure equidistant points to the left and right of the sagittal suture midline. A hole was drilled (Stereotaxic Drill, Kopf) at the coordinates corresponding to the target region. The dura was opened using a 25 G cannula, and the hole and skull surface were cleaned and kept moist with 0.9% isotonic saline (Fresenius Kabi) to prevent desiccation.

### Anterograde and retrograde viral injection

A syringe (Neuros™ Syringe; Model 7001; Hamilton) was loaded with virus prior to injection and positioned in the injector (Quintessential Stereotaxic Injector; Stoelting). The tip was positioned at the level of the dura and then slowly lowered manually to the target depth. After 2 minutes, virus was injected at a rate of 20 nl/min. After an additional 10 minutes, the syringe was slowly withdrawn manually. Following surgery, Betaisodona (Mundipharma GmbH) was applied to the suture site, and animals received a subcutaneous injection of Carprofen (5 mg/kg). Carprofen treatment (5 mg/kg, s.c.) was continued once daily for an additional three days.

Stereotaxic coordinates ^61^: CnF (AP: −7.8 mm, ML: +2.0 mm, DV: −5.8 mm), CeA (AP: −2.4 mm, ML: +4.2 mm, DV: −8.1 mm). Viral vectors and injected volumes: CnF (6.06 × 10^13^ gc/ml, 200 nl / 70 nl); central amygdala (1.35 × 10^14^ gc/ml, 35 nl).

### Microelectrode implantation

Five stainless steel screws (M1.6; length, 3 mm; Hummer & Rieß, Nuremberg, Germany) were inserted in boreholes without penetrating the dura overlaying the brain surface. Screws were inserted into each of the five holes. A platinum/iridium microelectrode (catalogue# UEPSEGSECN1M; FHC Inc., Bowdoin, ME, USA), with an impedance of 0.8-1.1 MΩ, was implanted to the coordinates: AP: −7.80 mm, ML: +2.00 mm, DV: −5.80, as described earlier ^20^. A custom-made plug (GT-Labortechnik, Arnstein, Germany) was attached to the electrode pin, and its ground wire was connected to one of the skull screws. The electrode and plug were secured to the skull screws using dental cement, which was applied to the skull and molded around the assembly to form a small protective cap ^68^. The skin anterior and posterior to the dental cement cap was sutured, and Betaisodona (Mundipharma GmbH) was applied. Animals received a subcutaneous injection of Carprofen (5 mg/kg). Postoperative analgesia was continued with Carprofen (5 mg/kg, s.c.) for an additional 3 days.

### Collection and processing of cerebral tissue

Once *in vivo* experiments were completed, rats were weighed and anaesthetised with an intraperitoneal injection of ketamine (90 mg/kg BW) and xylazine (10 mg/kg BW). After confirming loss of the hind-limb reflex, the thoracic cavity was opened and the heart exposed. A perfusion needle was inserted into the left ventricle and a small incision made in the right atrium. Rats were perfused with 1× PBS–0.4 % sodium-heparin (100 ml per rat) until the blood was cleared from the body, followed by fixation with 4 % paraformaldehyde (PBS-buffered, pH 7.4, 200 ml per rat). Brains were harvested and post-fixed in 4% paraformaldehyde for 1 day, washed in 1× PBS for 1 day, cryoprotected in 15 % and 30 % sucrose (1 day and 6 days, respectively), and embedded in Tissue-Tek O.C.T. Compound for cryosectioning.

Brain tissue was sectioned at 40 µm (coronal) or 60 µm (sagittal) at −21 °C using a CM3050 S cryostat (Leica). Sections were collected free-floating in 12- and 24-well plates (Nunclon™; Thermo Scientific) filled with 1× PBS and stored at 4 °C until immunohistochemical staining. For long-term storage, slices were preserved in a cryoprotection solution for free-floating sections (vol/vol; 30 % ethylene glycol, 20 % glycerol, 10 % 0.2 M phosphate buffer (pH 7.4), ∼ 40 % water.

### Immunohistochemistry

Cryosections were transferred to 4-well plates (Nunc multi dish 4-well, non-treated) and washed three times with 1× PBS. In case of c-Fos staining, sections were quenched with 100 mM glycine (pH adjusted to 7.4 with 2 M Tris base) and then permeabilised and blocked using a blocking solution (1× PBS; 10 % horse serum; 0.3 % Triton X-100; 0.1 % Tween 20), with each step performed for 30 minutes at room temperature. For tracing experiments, permeabilisation and blocking were performed for 60 minutes at room temperature, without a preceding quenching step.

Primary antibody incubation in blocking solution was carried out overnight at 4 °C (see Supplementary Table 2 for all primary and secondary antibodies). Sections were washed three times for 5 minutes each in wash buffer (1× PBS; 0.1 % Triton X-100; 0.1 % Tween 20) and then incubated with secondary antibody diluted in blocking solution for 1.5 hours at room temperature. Sections were washed four times for 10 minutes each in wash buffer, followed by a 10-minute incubation in 1× PBS with DAPI (2 mg/ml; 1:5,000) and washed again for 5 minutes. Stained sections were mounted on Superfrost Plus microscope slides (Avantor; VWR) and embedded in Aqua Poly/Mount (Polysciences).

### Microscopy

Stained cryosections were imaged using an Axio Imager M2 microscope (Zeiss) with ZEN 2.3 software, a 10x EC Plan-Neofluar air objective (0.3 NA) and an Axiocam 506 mono camera. Images were captured at excitation wavelengths of 353, 493, 545 and 650 nm. Binning was set to 4×4 for viral tracing and 2×2 for c-Fos imaging. Images comprised 91-170 tiles and were stored in 14-bit .czi format. Shading correction was applied to all acquired images.

For high-resolution imaging, an inverted Olympus IX81 microscope (Olympus) equipped with Fluoview FV10-ASW software, an FV1000 confocal laser scanning system, and a 60x UPLSAPO oil objective (1.35 NA) was used. The pinhole was set to one Airy unit. Images were acquired at excitation wavelengths of 405, 473, 559 and 635 nm, consisting of 14-19 z-layers (step size: 300 nm) and stored as 12-bit .oib files.

Images were processed using ImageJ (Fiji distribution; Wayne Rasband, National Institutes of Health, USA). Confocal images were processed as z-stacks per channel by maximum intensity projection. Brightness and contrast were adjusted, and merged channel composites created in RGB .tif format. The rat brain atlas ^61^ was used to identify different brain regions. Overlays were created with Adobe Illustrator and final image arrangements were performed in Adobe Photoshop (Adobe Systems Incorporated).

### Electrical deep brain stimulation of the mesencephalic locomotor region

Deep brain stimulation (cathodal; 60 µs pulse length; monophasic square-wave pulses) was applied using a cable-bound stimulus generator controlled by MC_Stimulus II software (Multi Channel Systems MCS GmbH). Stimulation began at 20 µA and was increased in 10 µA steps, each applied for 30 seconds (130 Hz), to determine the lowest behaviour-evoking amplitude (current threshold). This amplitude was then used to assess the frequency threshold for behavioural responses using 30-second stimulation bouts at 20, 30, 40, 50, 60, 80, 100, and 130 Hz.

#### Home cage stimulation for c-Fos *de novo* synthesis

Each rat was attached to the stimulation cable and placed in its home cage. To initially test stimulation conditions in the home cage, DBS was applied in 1-minute bouts, and behaviour was video-recorded with a smartphone camera. Rats then received high-frequency stimulation (130 Hz, 60 µs), low-frequency stimulation (40 Hz, 60 µs), or sham stimulation (cable attached, no current applied) for one hour, followed by a 30-minute resting period without the cable attached. The previously determined current threshold was used as the stimulation amplitude. Animals were subsequently sacrificed and their brains harvested.

#### Stimulation in an inverse open arena

Each rat was attached to the stimulation cable and placed in a transparent rectangular open field arena (80 × 42.5 cm). Animals were allowed to explore the arena freely before stimulation was applied for at least 30 seconds per condition. Behaviour was recorded from below (Basler ace Classic GigE colour acA1300-60gc camera with a 1/1.8” sensor and 4.4–11 mm/F1.6 Kowa varifocal lens) and from the side with a Sony FDR-AX700 4K camcorder. Simultaneously, real-time behavioural observations were documented manually to complement the video-based evaluation. Videos were acquired at frame rates of 40 and 25 fps. Before further processing, all DLC-derived coordinate-time series from the RCNN were curated and linearly interpolated to a common 40 fps grid.

### Pose estimation

For body part keypoint tracking and pose estimation, we used DeepLabCut (DLC, version 2.3.1) ^69^. Specifically, we trained a recurrent convolutional neural network (RCNN) on bottom-view videos of the rat open field. We utilized the pre-trained RCNN ‘*superanimal_topviewmouse*’ and labelled an additional *n* = 764 frames taken from *n* = 53 videos (then 95% was used for training). We iteratively trained a ResNet-152-based neural network with default parameters. The final model was trained with 900,000 iterations. Validation was performed using one shuffle; the test error was 6.67 pixels and the training error was 3.55 pixels (image size ≈ 1280 × 1024 pixels). A p-cutoff of 0.6 was used. This network was then used to analyse videos from similar experimental settings. Coordinate time series were further processed using custom-written MATLAB (R2022b, MathWorks, Natick, MA, USA) and Python (version 3.9 or 3.10) pipelines.

### Kinematics extraction from pose estimation data

Marker-based kinematics were extracted from *DeepLabCut* (DLC) tracking outputs using a custom MATLAB analysis pipeline. For each recording, pose-estimation files generated with the above described RCNN were identified from the project directory structure and organised into a metadata table according to the experimental conventions encoded in the file hierarchy. Tracking data were analysed at 40 frames/s. Only pose estimates passing a DLC likelihood cutoff of 0.99 were submitted to downstream kinematic extraction. Kinematic variables were computed using sliding temporal windows of 0.5 s.

Locomotor state was defined from the speed of the body centre marker. Frames were classified as locomotor behaviour when the rat centre speed exceeded 10 pixels s⁻¹. The following kinematic parameters were calculated from the tracked body-part coordinates: General locomotion was quantified from the rat centre as total distance travelled, maximum speed, minimum speed, mean speed, and median speed.

Speed summaries were calculated both across the full recording and restricted to locomotor epochs. In total, the extracted readouts comprised total distance travelled, rat-centre speed. For dynamic variables, summary statistics included extrema, means, medians, locomotion-restricted means and medians, as well as respective error measures.

### Immediate response kinematics curves

Open-field locomotion was quantified from *DeepLabCut* tracking of the “center” body point acquired at 40 frames/s. Coordinates with a *DeepLabCut* likelihood below 0.90 were treated as missing, and internal gaps of up to 0.5 s were linearly interpolated before speed calculation. Frame-to-frame Euclidean displacement was converted to speed in pixels per second, and values exceeding 1,000 px/s were excluded as tracking artefacts. Speed traces were summarized in consecutive non-overlapping 0.5-s bins as mean speed, median speed, maximum speed, and mean speed during locomotion, with locomotion defined as speed ≥ 10 px/s. Short internal gaps of up to 2 s in the resulting binned curves were interpolated without extrapolation at the beginning or end of recordings.

Repeated recordings were first averaged within each animal and experimental condition, such that animals rather than individual recordings constituted the statistical units. Analyses compared Off with HFS and LFS conditions. Curves were restricted to the duration shared by all animal–condition datasets included in the respective comparison and are presented as animal-level mean ± s.e.m.

For statistical analysis, each animal’s mean value was calculated over the first 5 s and first 10 s of each recording. Animals were included only when at least 80% of the expected time bins were available in both Off and the respective stimulation condition. Conditions were compared with Off using two-sided paired Wilcoxon signed-rank tests. Holm correction was applied separately across the condition-versus-Off comparisons within each metric and time interval.

### Semi-supervised behavioural classification

For more controlled capture of predefined behavioural motifs, and downstream unsupervised discovery of sub-behaviours within those motifs ^70^, we used a semi-supervised machine learning approach, the *A-SoiD* pipeline ^47^. Specific behaviours were annotated in an ethogram using *BORIS* software ^71^, combined with corresponding pose estimation data from *DeepLabCut* ^72^. We extensively trained an active learning classifier on a diverse yet balanced set of bottom-view pose estimation data from a total of 34 videos, analysed within DLC using the model described above. Each frame of every training video (*n* = 34) was then annotated in *BORIS* with one of 10 behavioural categories: *Circling*, *Fast locomotion–Running*, *Freezing*, *Grooming*, *Idle-Exploration*, *Immobility*, *Rearing*, *Slow locomotion–Walk*, *Tail rattling*, and *Turning*. Ethograms were exported as binary behaviour files at 40 Hz.

Training (hyper)parameters for the A-SoiD active learning classifier were set as follows: DLC likelihood cutoff, 0.94; minimum bout duration, 0.05 s; training fraction, 0.03; test fraction, 0.97; maximum number of iterations, 1,000; confidence threshold for inclusion in the training dataset, 0.5; video frame rate, 40 fps; ethogram rate, 40 Hz.

The final *A-SoiD* classifier (random forest) achieved moderate-to-high average performance for most classes (≥ 0.80 F1 score), with “*Turning*” the poorest-performing category (∼ 0.60 F1 score).

### A-SoiD: Behavioural analysis

For each recording, frame-wise behavioural labels were loaded from the corresponding .npy file. Metadata were parsed from the filename, including stimulation state, behavioural sub-condition, animal identifier, trial, stimulation amplitude, amplitude group, stimulation frequency, frequency group, and repetition. Behavioural class names and the recording frame rate were read from the *A-SoiD* configuration file. For stimulation-off recordings, frequency and amplitude labels were recoded as “Off” to allow direct comparison with stimulation-on conditions. For each recording, behavioural bouts were defined as consecutive frames assigned to the same *A-SoiD* class. Bout duration was calculated as bout length divided by the video frame rate. For each animal, behavioural class, and experimental condition, we quantified mean bout duration, bout count, and the fraction of recording time spent in each behaviour. Metrics were first averaged across repetitions and then averaged at the animal level for each condition. Behavioural classes labelled “other” were excluded from downstream statistical analyses and visualisation.

Transition probabilities were computed from frame-to-frame changes in the *A-SoiD* label sequence. For each recording, a directed transition matrix was constructed across all behavioural classes and row-normalised to yield the probability of transitioning from each source behaviour to each target behaviour. Transition matrices were then averaged across repetitions and animals using the same condition structure as for bout metrics.

#### Statistical analysis of behavioural metrics

Statistical analyses were performed separately for each behavioural class and experimental factor. For bout counts and mean bout durations, animal-level values were arranged as paired repeated-measures tables across factor levels. Where two levels were present, paired comparisons were performed using two-sided Wilcoxon signed-rank tests. For factors with more than two levels, Friedman tests were used as omnibus repeated-measures tests; where these were significant, post hoc comparisons were performed using Nemenyi tests when available, or otherwise pairwise Wilcoxon signed-rank tests with Holm correction. Omnibus P values were additionally corrected using the Benjamini–Hochberg false-discovery-rate procedure. Statistical significance was assessed at α = 0.05.

Bout-duration distributions were compared between conditions using empirical cumulative distribution functions. For each behavioural class, multi-group distributional differences were assessed using Anderson–Darling k-sample tests, and pairwise distributional comparisons were performed using two-sample Kolmogorov–Smirnov tests with multiplicity-adjusted P values.

#### Behavioural correlation and temporal coupling analyses

To assess relationships between behavioural classes, animal-level bout counts were pivoted into behaviour-by-animal matrices for each condition, and Pearson correlation coefficients were calculated between behavioural classes within conditions. Between-condition behavioural correlation matrices were additionally computed by correlating behaviour counts across matched animals between pairs of conditions. Resulting P values were corrected using the Benjamini–Hochberg false-discovery-rate procedure.

Temporal coupling between behavioural classes was quantified using frame-resolved binary vectors for each *A-SoiD* class. For each pair of behaviours, binary label vectors were z-scored and cross-correlated over a maximum lag of ± 5 s; the maximal absolute normalised cross-correlation and its corresponding lag were extracted for each recording and averaged per animal. Between-condition differences in maximal cross-correlation and lag were assessed using paired Wilcoxon signed-rank tests across animals, followed by Benjamini–Hochberg correction across behaviour pairs.

#### Low-dimensional embedding and visualization

To visualise behavioural structure across stimulation-related experimental factors, we generated three-dimensional nonlinear embeddings from averaged behavioural feature tables using UMAP and t-SNE. The input table contained per-animal or per-condition behavioural summaries, including behaviour identity, time proportion, mean bout duration, and bout count. Analyses were performed separately for predefined experimental factors (Off vs. HFS; LFS had too little values for reliable embedding). For each factor level, rows with missing values in the selected metric, TimeProp, or behaviour label were excluded, as were rows labelled “Other“; factor levels labelled “None” were omitted. Embeddings were computed only for subsets containing at least ten observations.

For each subset, behaviour labels were one-hot encoded and combined with either mean bout duration or bout count together with time proportion. In a second feature-space variant, transition-matrix features were additionally merged from the corresponding transition summary table. Transition features could contain multiple rows per animal and factor level. Therefore, transition matrices were first averaged within each Animal ID × factor combination before merging, to prevent many-to-many duplication. Features containing only missing values or no variance were removed, infinite values were treated as missing, and remaining missing values were mean-imputed.

For each factor level and metric, three-dimensional t-SNE embeddings were computed using perplexities of 10, 30, and 50 (restricted to values smaller than the number of observations in the analysed subset), also with a fixed random seed of 42. Embedding coordinates were plotted in three dimensions and coloured by behaviour identity using a viridis colour scale.

### Un-supervised behavioural classification

We trained a full, location-aware *Keypoint-MoSeq* (KPMS) model (a switching linear dynamics system (SLDS) model) as described previously ^48^ after first fitting an initial autoregressive Hidden Markov Model (AR-HMM) with a kappa (stickiness) hyperparameter of 10^13^ (dimensionless), yielding a median syllable duration of ∼ 20 ms. Since coordinate time series were sampled at 40 fps and the median syllable duration was ∼ 20 ms in AR-HMM terms, syllables were on average ∼ 800 ms long. Principal component analysis (PCA) on the AR-HMM output showed that 10 principal components explained ≥ 90% of the variance. The kappa hyperparameter was maintained at 10^13^ when fitting the full model (final median syllable duration at 350 ms). The model was applied to all *n* = 810 bottom-view open field videos from *n* = 22 rats, yielding 19 principal behavioural syllables (syllable 0 → 18 = decreasing order of frequency of occurrence), of which syllables with a frequency > 0.005 were retained for analysis (see the full, non-restricted syllable set in Supplementary Figure 3).

Downstream statistical analysis and plotting was conducted as described previously ^48, 73^, with modifications described below. KPMS outputs were analysed using a custom Python workflow. Per-recording result files were first organised by experimental condition using filename-derived stimulation metadata; recordings were assigned to Off, weak high-frequency stimulation (HFS), strong HFS, weak low-frequency stimulation (LFS), or strong LFS groups according to the stimulation amplitude– frequency combination encoded in the file name. For the animal-based analysis, KPMS summary tables were loaded from stats_df.csv files, and animal identity was extracted from the recording name to generate a subject_id. All subsequent statistical analyses were performed at the animal level, by averaging each behavioural metric across group, animal, and syllable. Syllable-level analyses were restricted to the predefined set of model syllables used for downstream interpretation, with rare syllables routinely excluded using a minimum frequency threshold of 0.005.

For each retained syllable, the following KPMS-derived readouts were analysed: syllable frequency; syllable duration; mean, standard deviation, minimum, and maximum heading; mean, standard deviation, minimum, and maximum angular velocity; and mean, standard deviation, minimum, and maximum linear velocity (pixels/s). Syllable names were mapped to manually curated behavioural labels using trajectory and grid movies and used via the model’s syllable_names.csv file. These labels were used in figure titles and axis annotations.

For each metric and syllable, group differences were tested only when at least two groups contained data from three or more animals. Non-parametric statistics were routinely used, as syllable usage and duration distributions were not assumed to be normally distributed. Group effects were assessed with Kruskal–Wallis tests applied to animal-averaged values. P values were estimated using 10,000 blocked permutations, preserving the within-animal structure of the data, and were corrected across syllables using the Benjamini–Hochberg false-discovery-rate procedure. When the Kruskal– Wallis test was significant, predefined pairwise comparisons were evaluated using Dunn’s post hoc tests. The planned contrasts were Off vs. strong HFS, Off vs. weak HFS, Off vs. strong LFS, and weak HFS vs. strong HFS (weak LFS excluded due to low sample size). Statistical significance was defined as adjusted *P* < 0.05. Results of the Kruskal–Wallis and post hoc analyses were exported as per-syllable CSV files.

To analyse behavioural sequencing, transition matrices were generated from the frame-wise KPMS syllable labels. Consecutive repeats of the same syllable were collapsed so that transitions reflected changes between behavioural states rather than dwell time within a state. Bigram transition matrices were computed for each recording and combined within each experimental group, then normalised to yield transition probabilities; only syllables exceeding the frequency threshold were retained. Group-level transition matrices were visualised as heat maps, with rows and columns corresponding to incoming and outgoing syllables, respectively. To allow direct comparison across groups, heat maps used a common colour scale defined by the maximum transition probability observed across group matrices.

Differences in transition structure between experimental groups were visualised by subtracting one group-level transition matrix from another for predefined comparisons; these difference matrices were plotted as heat maps to identify transitions enriched or depleted under stimulation relative to Off, or between stimulation strengths. In parallel, transition networks were generated using transition probabilities as edge weights, with syllables represented as nodes (node size scaled by syllable usage) and edges scaled by transition probability. Difference networks were also plotted, with node and edge colour indicating the direction of change between groups.

Finally, global differences in transition architecture were assessed using spectral analysis of the group-level transition matrices. Eigenvalue spectra were computed for each matrix, and pairwise differences between groups were quantified using the Frobenius norm of the eigenvalue differences. Statistical significance of spectral differences was estimated by permutation testing (1,000 permutations). Resulting P values were Bonferroni-corrected for the number of predefined group comparisons.

For KPMS syllable cluster analysis, syllables were pooled according to a trajectory-similarity dendrogram using a clustering scheme. The four-cluster scheme comprised syllable 15 = cluster 1; syllables 10, 13, 3 and 9 = cluster 2; syllables 5, 17, 8, 14, 7, 0, 1, 16, 12, 2 and 6 = cluster 3; and syllables 11, 4 and 18 = cluster 4. Only syllables with a global frame-occupancy frequency > 0.005 were included, with this cutoff applied before syllables were pooled into clusters. For each recording, relative cluster usage was calculated as the fraction of retained, cluster-assigned frames belonging to the respective cluster. Mean in-cluster speed was calculated as the frame-weighted mean of velocity_px_s, and cluster duration was calculated as the mean duration of the original constituent-syllable bouts at 40 frames/s. Transitions between different syllables assigned to the same cluster were retained as separate bouts. Analyses were conducted for Off, HFS and LFS, in which weak and strong stimulation levels were pooled. Repeated recordings were first averaged within each animal and condition, such that animals constituted the statistical units. For each cluster and outcome measure, stimulation conditions were compared with Off using two-sided paired Wilcoxon signed-rank tests, including only animals with valid measurements in both conditions and requiring at least three matched animals. Raw *p* values were Holm-corrected across all condition-versus-Off comparisons displayed in the same violin plot, encompassing all clusters and stimulation conditions for the respective metric and condition grouping.

### Deep learning-based quantitative image analysis for c-Fos

Image segmentation was conducted using *deepflash2* ^51^, following established guidelines for reproducible bioimage analysis ^50^. The c-Fos-channel was extracted from the Zeiss .czi files and converted to .tiff file format using Python. In *QuPath* ^74^, three experts annotated six images with five regions of interest (ROIs) each. The annotated cFos coordinates were exported using a *QuPath* macro and converted to binary masks in Python. The simultaneous truth and performance level estimation (STAPLE) method was used for ground truth estimation (provided in *deepflash2* GUI; Supplementary Table 1). The images were randomly divided into 80% training and 20% testing data, and a model ensemble was formed by training three consensus deep learning models. The Dice score was used to assess model accuracy against test set annotations (Supplementary Table 1). The trained model ensemble was used to segment c-Fos images, with segmentation quality assessed based on test set performance and visual inspection. Images from four LFS animals required brightness and contrast correction.

For the area-specific quantification of cFOS+ cells/mm², four regions were defined according to the Paxinos and Watson rat brain atlas ^61^: “CnF”, including the dorsal, intermediate and ventral part of the CnF and the precuneiform nucleus; “PPN”, including the pedunculopontine nucleus; “CIC”, including the dorsal cortex, external cortex and central inferior colliculus; “PAG”, including the dorsolateral, dorsomedial, lateral, ventrolateral periaqueductal gray, laterodorsal tegmental nucleus, the dorsal and ventral part of the dorsal raphe nucleus and the posterodorsal raphe nucleus. Each region was manually annotated on the ipsilateral and contralateral side in *QuPath*, resulting in eight defined regions. The annotations were exported with a groovy script as .json files and further processed in Python. Region coordinates were converted to binary masks, overlaid with the corresponding predicted cFos binary masks, and the area, cFos+ cell count, and the cFos+ cells/mm^2^ were calculated for each region.

### Statistical analysis, plotting and data reporting

Statistical analyses and plotting of analyses were conducted using GraphPad Prism (Version 10.0) or custom-built MATLAB (R2022b, MathWorks, Natick, MA, USA) and Python (Python, Version 3.9 or 3.10) scripts. Assessment for normality was conducted by using normality/lognormality testing (most reliable test according to sample size was used) and assessment of Q-Q plots. Individual statistical testing was conducted on a group level via analysis of variance (ANOVA, one- or two-way) or mixed-effects models whenever possible and relevant, to assess for factor and interaction effects. In case of non-gaussian distribution of data, corresponding standard non-parametric equivalents were used. Source data and custom code is available from the corresponding authors upon reasonable request. Significance is annotated as ns = p>0.05, *p<0.05, **p<0.01, ***p<0.001, ****p<0.0001, if not stated otherwise. Figures were created in Adobe Illustrator. Publicly available sketches from SciDraw were used.

## Supporting information

supplemental video 1: DBS in the CUN of the rat-behavior

## Data availability

Requests for further information and resources should be directed to and will be fulfilled by the lead contact, Robert Blum.

## Code availability

Custom codes are available from the corresponding author upon request.

## Acknowledgements

We thank Hildegard Troll for AAV virus production. This work was funded by the Deutsche Forschungsgemeinschaft (DFG, German Research Foundation), project ID: 424778381 (CRC Retune, A01, A02, B06) and the Interdisziplinäres Zentrum für Klinische Forschung (IZKF) Würzburg, grant S-530.

## Author contributions

Conceptualization, M.K.S., J.V., R.B.; Methodology, J.S.G., F.S., A.S., F.F., R.B.; Formal Analysis: F.S., A.S., J.S.G., J.H.; Investigation, M.W., TCF, D.S., S.M., F.H., S.K., F.F., R.B.; Data Curation, M.W., J.H., D.S., S.M., F.H., R.B. Writing – Original Draft, M.W., J.H., and R.B.; Writing –Review & Editing, M.W., J.H., P.T., C.W.I., F.F., M.K.S., J.V., R.B. Supervision, P.T., M.K.S., R.B.; Funding Acquisition, P.T., C.W.I., M.K.S., J.V., R.B. Project Administration, J.V., R.B.

## Additional information

**Supplementary information** The online version contains supplementary material.

### Supplementary material

**Supplementary Fig. 1.**
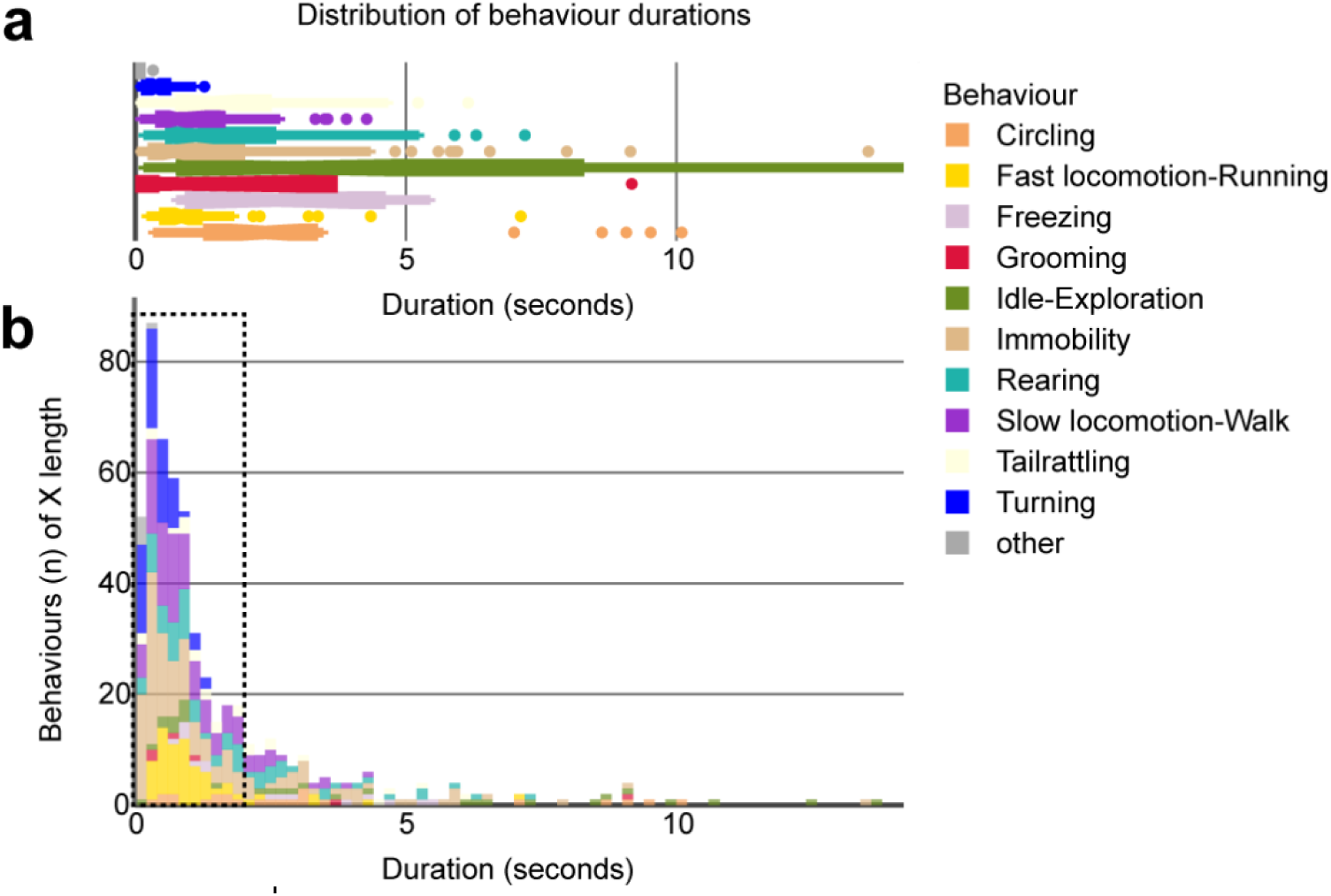
Duration of captured behavioural categories by semi-supervised machine learning are mostly < 2 seconds. The plot shows relative bout counts (y-axis) of each annotated behaviour and its respective duration in seconds (x-axis). The dotted box encloses the most frequent behavioural bouts, which show lengths of mostly < 2 seconds.

**Supplementary Fig. 2.**
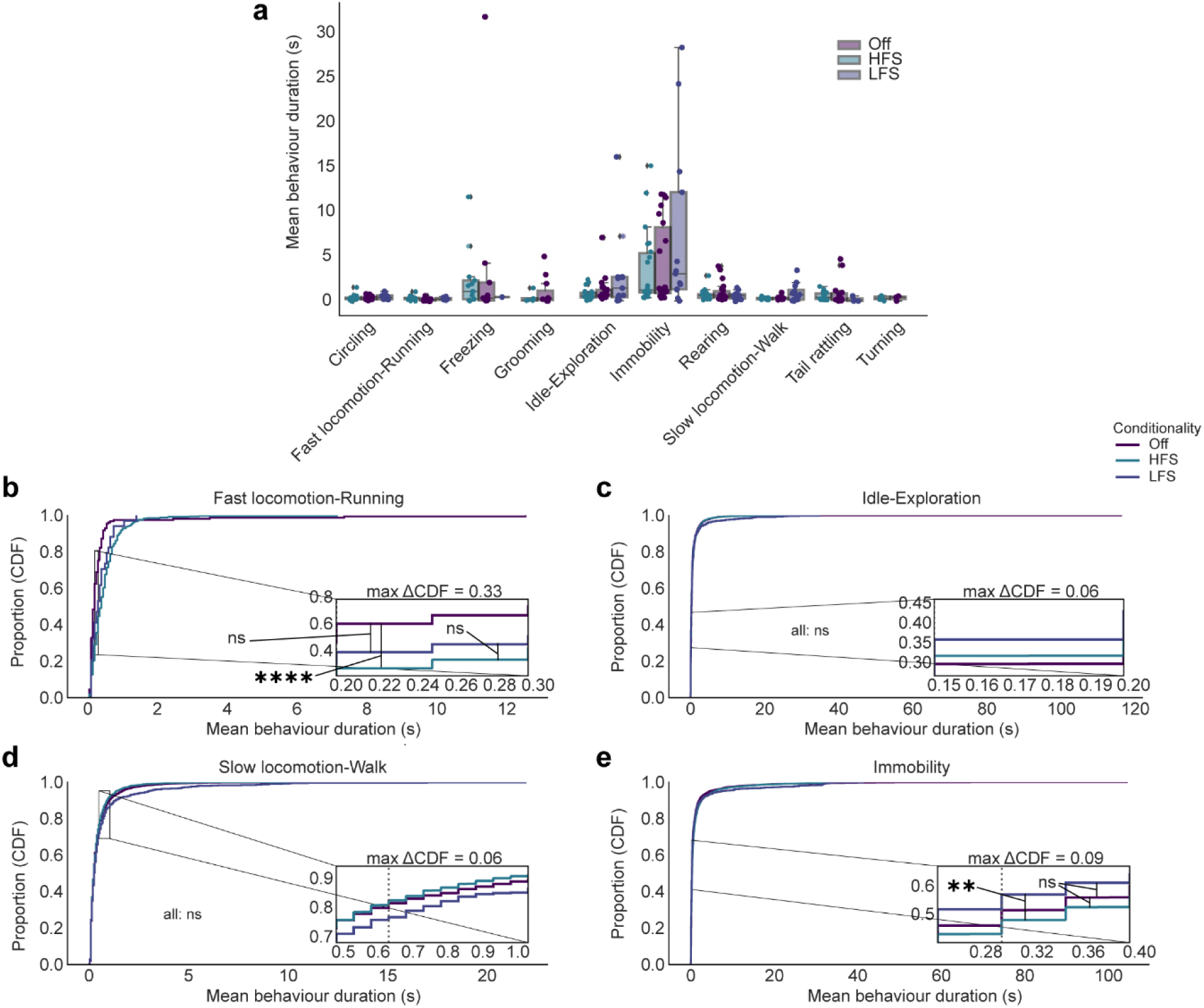
High-frequency MLR-DBS leads to proportionally longer running bouts. **a.** Mean bout durations of the different behaviours identified by the semi-supervised machine learning classifier across n = 810 videos showed no significant changes (one Friedman test per behaviour with Nemenyi post-hoc test). **b.** Cumulative distribution function (CDF) histograms display the proportion (i.e. fraction) of different length of bouts of behaviours. Comparison for each behaviour across conditionalities with multi-sample Anderson–Darling test with two-sample Kolmogorov–Smirnov tests with Bonferroni correction. Running (Statistic: 31.75, p = 0.001) bouts are prolonged via HFS (p < 0.0001), but not LFS (p = 1). **c.** Idle exploration behaviour (Anderson-Darling statistic: 4.666, p = 0.0026) bouts show changes, however all non-significant (all p > 0.05). d. Slow locomotion (Statistic: 3.514, p = 0.0091) bouts show changes, however all non-significant (all p > 0.05). **d.** Immobility bouts (Statistic: 14.69, p = 0.001) bouts show changes, with a significant different between HFS and LFS (p = 0.0010)

**Supplementary Fig. 3.**
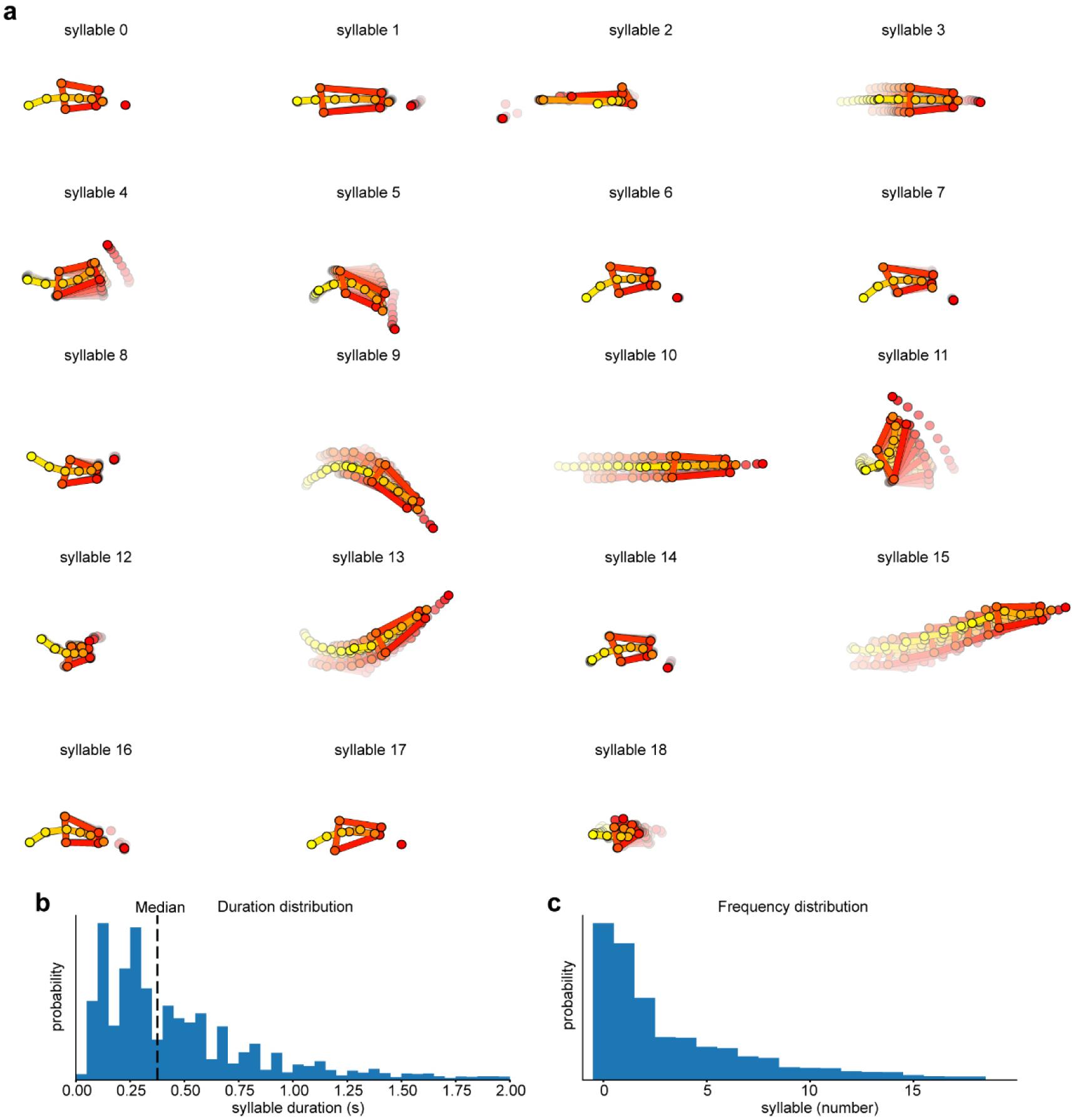
Unsupervised machine learning workflow and methodology via Keypoint-MoSeq. **a.** Syllable trajectories (n = 19) of behaviour identified by a trained Keypoint-MoSeq model. The classifier was trained on n = 810 videos from n = 22 rats. **b.** Plot showing the distribution of syllables of different duration (in seconds) according to probability of occurrence. Median syllable duration value at 350 ms (dotted line). **c.** Plot showing the distribution of different syllable identities with respect to probability of occurrence.

**Supplementary Fig. 4.**
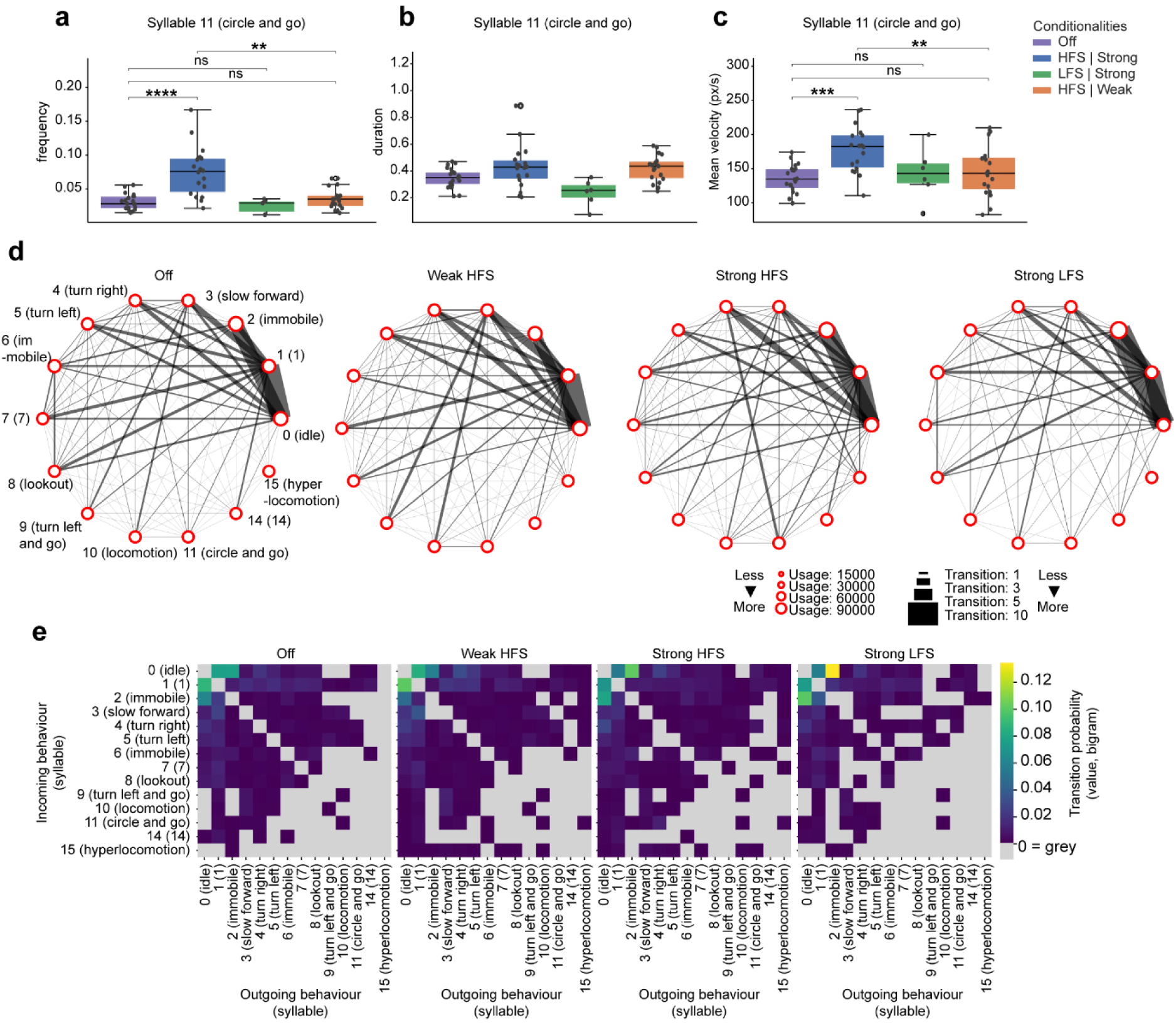
Changes of kinematic parameters induced by LFS vs. HFS. **a.** Frequency (Kruskal-Wallis test: H-statistic: 24.64, p = 0.0001) during a syllable displaying a turn behaviour followed by fast locomotion (syllable 11, Circle and go) is strongly increased by both high-amplitude HFS (p < 0.0001) but not low-amplitude HFS (p = 0.3871). **b.** Duration of syllable 11 (Kruskal-Wallis test: H-statistic: 14.03, p = 0.0026) is not changed by DBS-conditions. **c.** Mean velocity of syllable 11 (Kruskal-Wallis test: H-statistic: 14.40, p = 0.0005) shows increase weak and strong high-amplitude HFS (p = 0.0009), but not low-amplitude HFS (p = 0.2999). **d.** Native node and network plots of each conditionality showing usage (node size) and transition magnitudes (line thickness) per conditionality. **e.** Native transition matrices of different conditionalities, displaying the probability of transitioning from each syllable into the others. Strong LFS enhanced the probability of transitioning between idle exploration syllable and immobility syllable by ∼ 12%. Syllable statistics all as Kruskal-Wallis tests with animal-level clustered permutations and post-hoc Dunn‘s z-test (Holm-adj. sign. depicted) per syllable across groups. Box plots show median, IQR with whiskers as most extreme data points. Sample sizes: Off: n = 22; HFS: n = 22; LFS: n = 13, if not indicated otherwise.

**Supplementary Fig. 5.**
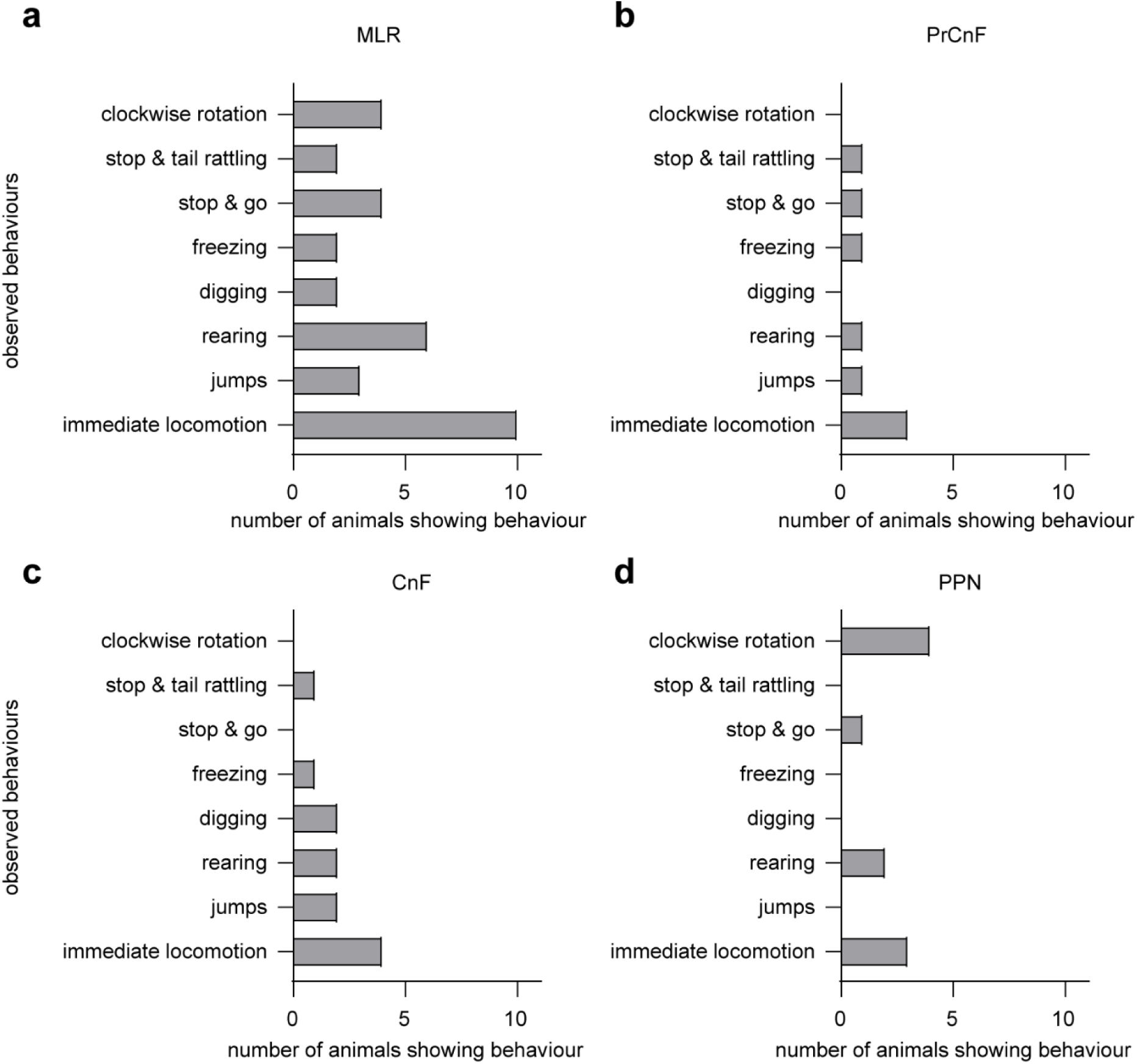
Observed, acute home-cage behaviours induced by high-frequency-stimulation in the MLR. Bars indicate the number of animals exhibiting the indicated behaviour after high-frequency-stimulation (HFS). Animals could show more than one of the behavioural categories or none. **a.** Behaviour of animals that received HFS-DBS in the MLR (n = 11). **b - d.** Behaviour of animals with electrode tip placement in the precuneiform nucleus (PrCnF) of the MLR, n = 3/11, in b.), cuneiform nucleus (CnF; n = 4/11, in c.), or pedunculopontine nucleus (PPN; n = 4/11, in d.).

**Supplementary Fig. 6.**
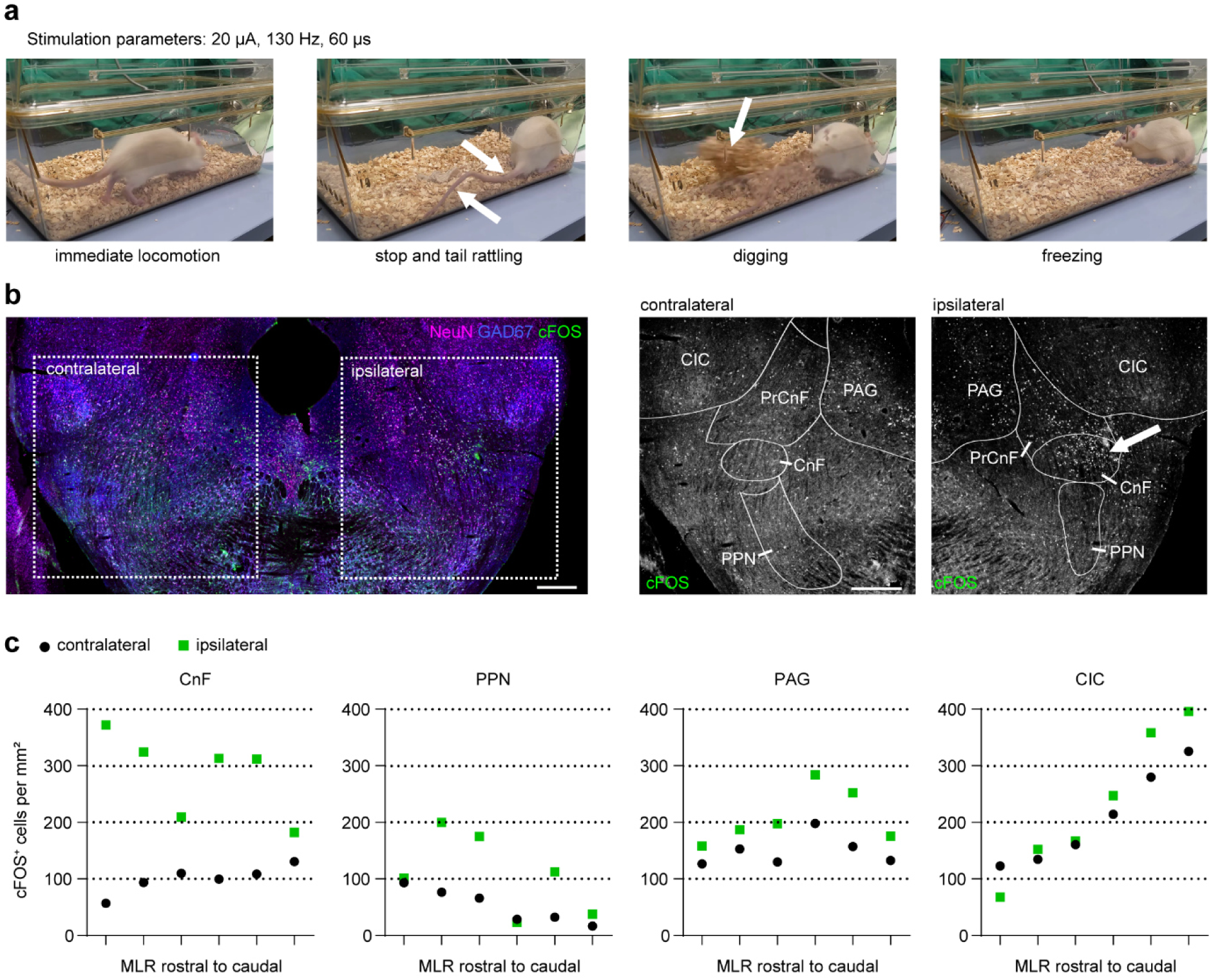
Single animal experiment example, rat with electrode placement in the CnF: observed behaviours, c-Fos distribution pattern induced by a 60 min DBS-period. **a.** Behaviours elicited by HF-DBS. Behaviours observed during stimulation at 20 µA, 130 Hz and 60 µs included immediate locomotion, stopping with tail rattling, digging, and freezing. Arrows indicated episodes of tail rattling and digging. **b.** Overview and regional distribution of c-Fos expression. Left: Coronal section of the mesencephalic locomotor region (MLR) showing c-Fos (green), NeuN (magenta), and GAD67 (blue) to outline GABAergic structures. Right: Higher-resolution views of contralateral and ipsilateral (stimulated) sides showing c-Fos labelling with anatomical overlays delineating the precuneiform nucleus (PrCnF), cuneiform nucleus (CnF), pedunculopontine nucleus (PPN), inferior colliculus (CIC) and periaqueductal grey (PAG). The arrow indicates the electrode position. Scales, 1 mm. **c.** Quantification of c-Fos-positive cells. Number of c-Fos-positive cells per mm² in the contralateral and ipsilateral CnF (including PrCnF), PPN, CIC and PAG. Data are shown for six coronal sections spanning the rostro-caudal extent of the MLR. The first contra- and ipsilateral data points represent the section shown in b.

**Supplementary Fig. 7.**
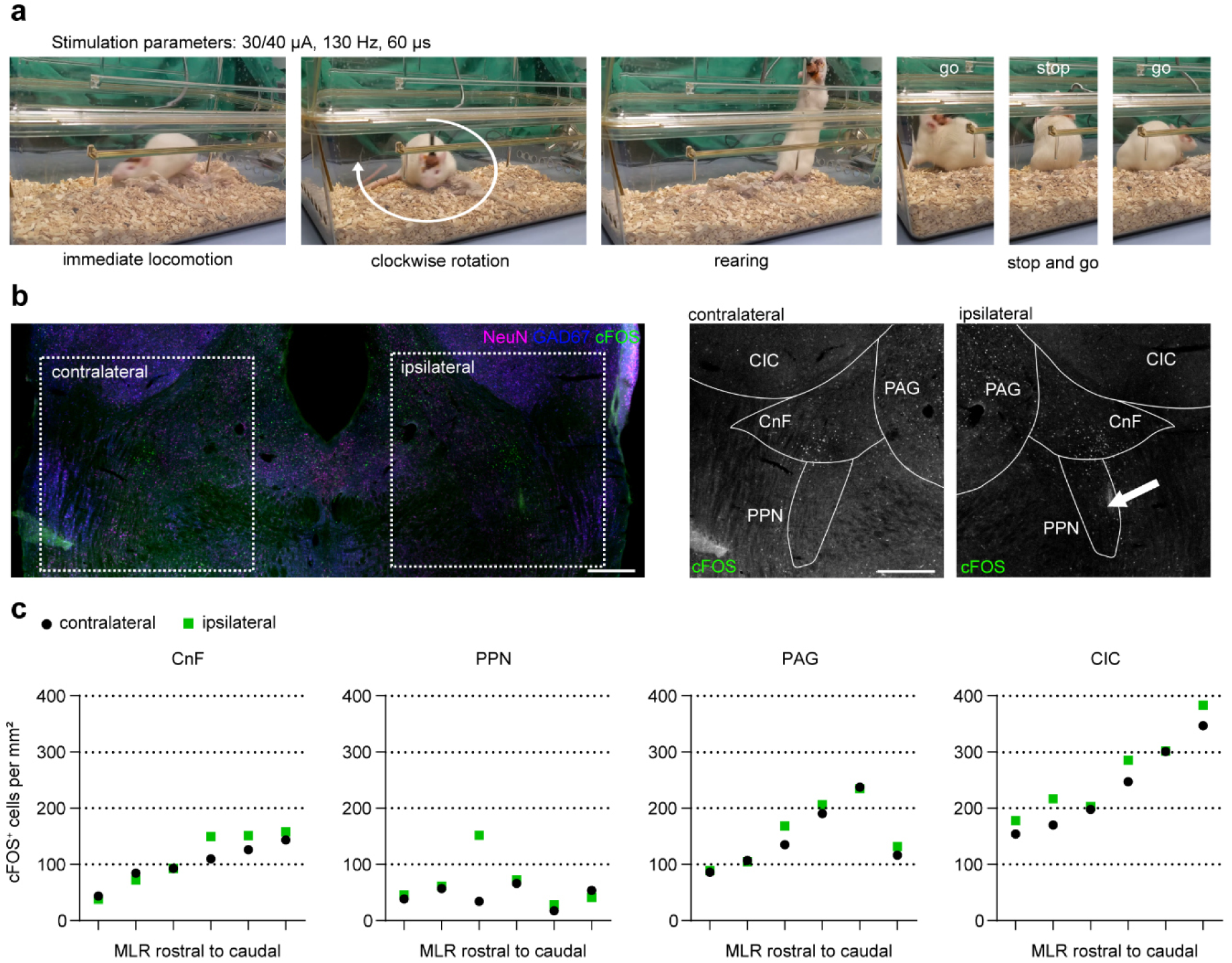
Single animal experiment example, rat with electrode placement in the PPN: observed behaviours, c-Fos distribution pattern induced by a 60 min DBS-period. **a.** Behaviours elicited by HF-DBS. Behaviours observed during stimulation included immediate locomotion at 40 µA (first image) and clockwise rotation, stop and go and rearing at 30 µA (130 Hz and 60 µs). **b.** Overview and regional distribution of c-Fos expression. Left: Coronal section of the mesencephalic locomotor region (MLR) showing c-Fos (green), NeuN (magenta), and GAD67 (blue) to outline GABAergic structures. Right: Higher-resolution views of the contralateral and ipsilateral (stimulated) sides showing c-Fos labelling with anatomical overlays delineating the cuneiform nucleus (CnF), pedunculopontine nucleus (PPN), inferior colliculus (CIC) and periaqueductal grey (PAG). The arrow indicates the electrode position. Scales, 1 mm. **c.** Quantification of c-Fos-positive cells. Number of c-Fos-positive cells per mm² in the contralateral and ipsilateral CnF, PPN, CIC and PAG. Data are shown for six coronal sections spanning the rostro-caudal extent of the MLR. The fourth contra- and ipsilateral data points represent the section shown in b.

**Supplementary Fig. 8.**
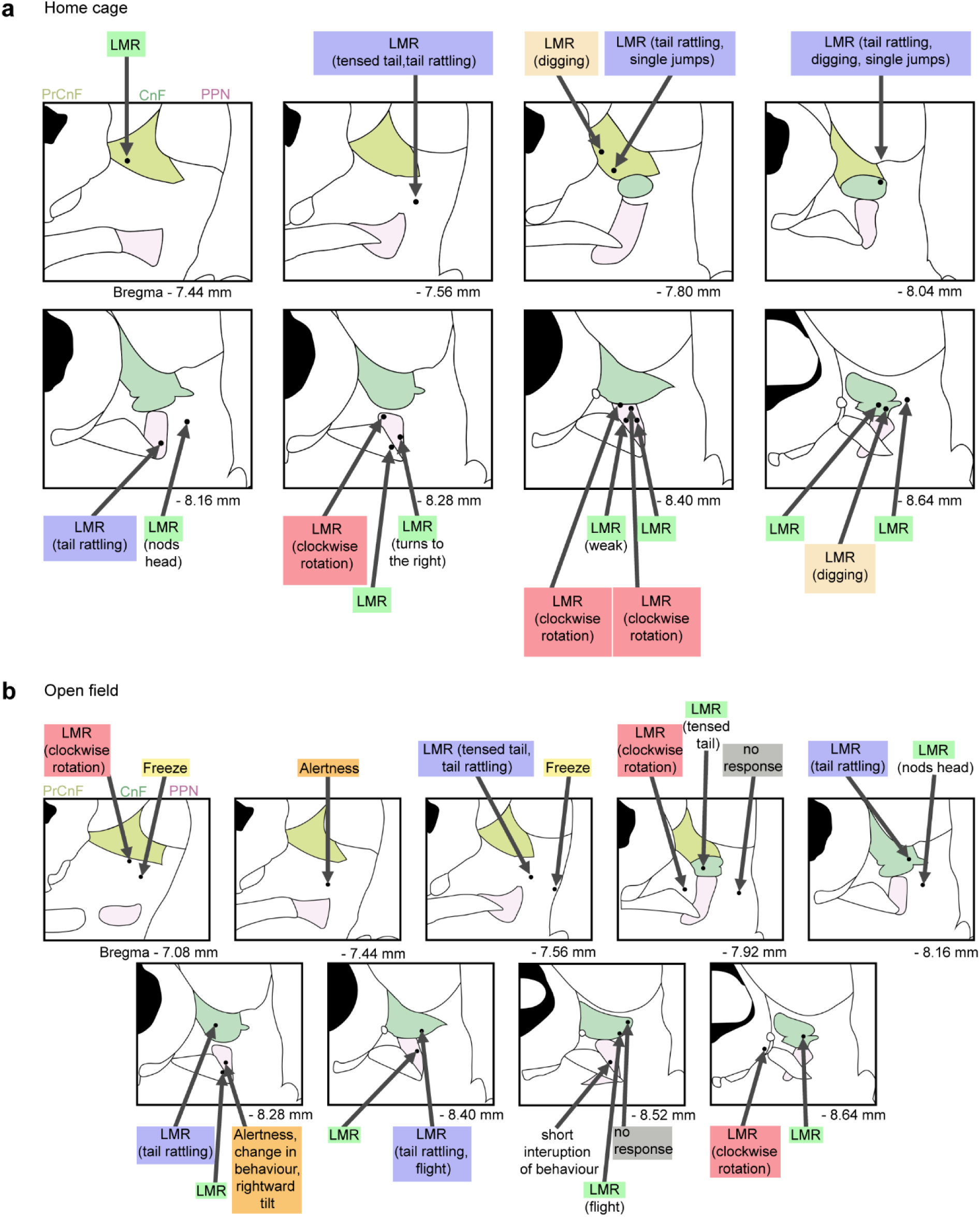
Stimulation electrode tip positions and subjectively observed acute DBS behaviours. DBS electrode tip positions are mapped onto outlines of the MLR and neighbouring regions according to coronal sections of the rat brain ^61^. Stereotactic coordinates are indicated relative to Bregma. Arrows indicate individual experiments and the acute behavioural responses observed during high-frequency, high-amplitude DBS. **a**. Observations during DBS in the home cage (c-Fos mapping experiment). **b.** Observations during DBS in the open arena.

### Supplementary Tables

**Supplementary Table 1.** c-Fos model scores.

| **Ground truth estimation scores of expert annotations** |  |  |
| --- | --- | --- |
| *expert* | *average_dice_score* | *std_dice_score* |
| 1 | 0.802 | 0.180 |
| 2 | 0.686 | 0.338 |
| 3 | 0.783 | 0.201 |
| **mean** | **0.757** | **0.240** |
| **Performance of c-Fos model ensemble on validation images** |  |  |
|  | *Mean dice score* | *Mean uncertainty score* |
| model 1 | 0.817 | 0.054 |
| model 2 | 0.755 | 0.050 |
| model 3 | 0.773 | 0.046 |
| **mean** | **0.782** | **0.050** |
| **Performance of c-Fos model ensemble on test images** |  |  |
| *file* | *dice_score* | *uncertainty_score* |
| 03_0.tif | 0.853 | 0.043 |
| 03_1.tif | 0.661 | 0.100 |
| 03_2.tif | 0.738 | 0.073 |
| 03_3.tif | 0.344 | 0.037 |
| 03_4.tif | 0.660 | 0.089 |
| **mean** | **0.651413** | **0.068** |

For train/test split: 5 image sections of 6 brain slices were annotated by three experts and the ground truth was estimated using the STAPLE method. 5 images of one brain slice were used for testing the model, the other images were used for training and validation of an ensemble model (Unet\_resnet34\_2classes, 3 models). Learning rate: 0.0005 (for all other parameters, the default parameters of deepflash2 (v0.1.8 ) was used).

**Supplementary Table 2.**
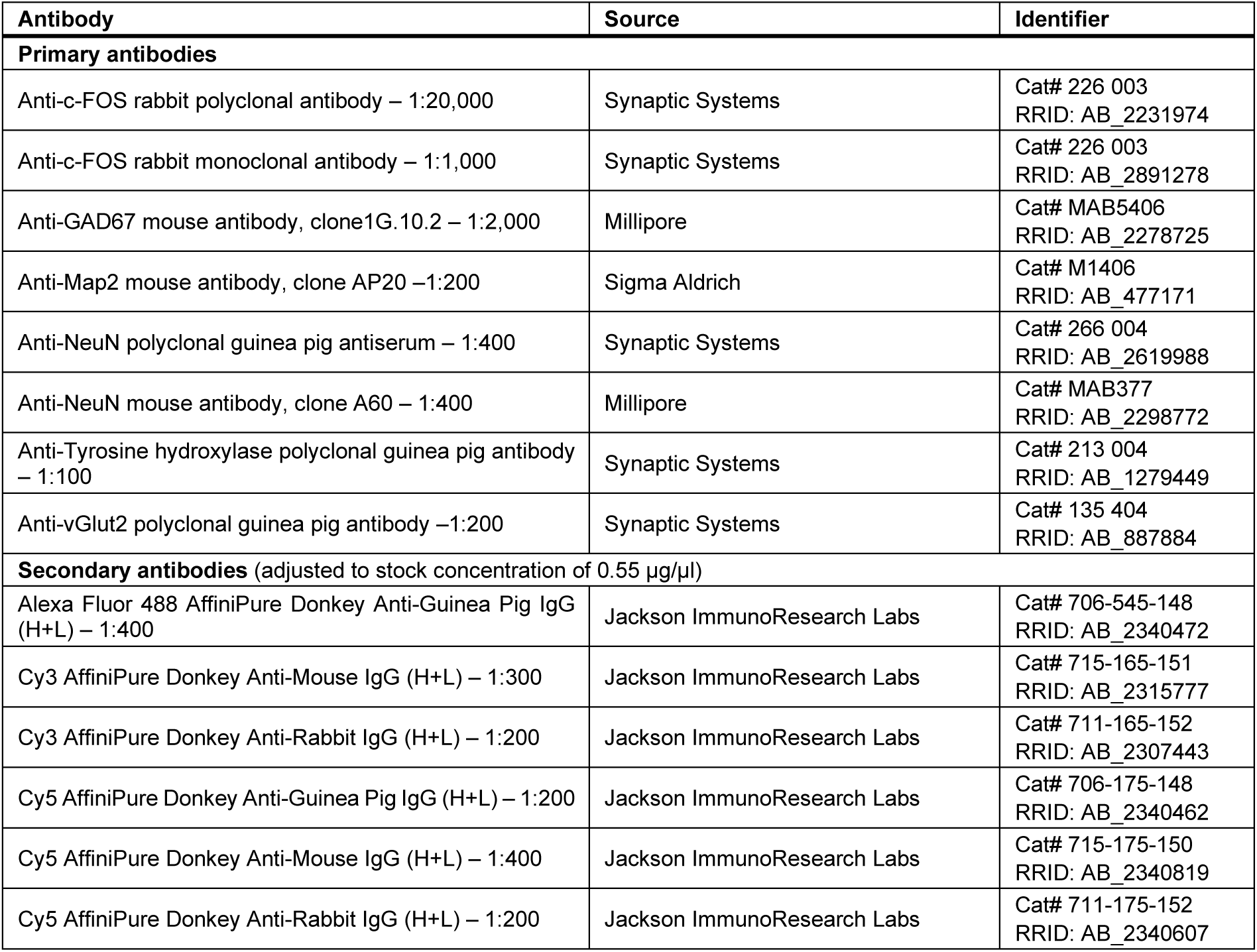
Primary and secondary antibodies used.

| Antibody | Source | Identifier |
| --- | --- | --- |
| **Primary antibodies** |  |  |
| Anti-c-FOS rabbit polyclonal antibody – 1:20,000 | Synaptic Systems | Cat# 226 003<br>RRID: AB_2231974 |
| Anti-c-FOS rabbit monoclonal antibody – 1:1,000 | Synaptic Systems | Cat# 226 003<br>RRID: AB_2891278 |
| Anti-GAD67 mouse antibody, clone 1G.10.2 – 1:2,000 | Millipore | Cat# MAB5406<br>RRID: AB_2278725 |
| Anti-Map2 mouse antibody, clone AP20 – 1:200 | Sigma Aldrich | Cat# M1406<br>RRID: AB_477171 |
| Anti-NeuN polyclonal guinea pig antiserum – 1:400 | Synaptic Systems | Cat# 266 004<br>RRID: AB_2619988 |
| Anti-NeuN mouse antibody, clone A60 – 1:400 | Millipore | Cat# MAB377<br>RRID: AB_2298772 |
| Anti-Tyrosine hydroxylase polyclonal guinea pig antibody – 1:100 | Synaptic Systems | Cat# 213 004<br>RRID: AB_1279449 |
| Anti-vGlut2 polyclonal guinea pig antibody – 1:200 | Synaptic Systems | Cat# 135 404<br>RRID: AB_887884 |
| **Secondary antibodies** (adjusted to stock concentration of 0.55 µg/µl) |  |  |
| Alexa Fluor 488 AffiniPure Donkey Anti-Guinea Pig IgG (H+L) – 1:400 | Jackson ImmunoResearch Labs | Cat# 706-545-148<br>RRID: AB_2340472 |
| Cy3 AffiniPure Donkey Anti-Mouse IgG (H+L) – 1:300 | Jackson ImmunoResearch Labs | Cat# 715-165-151<br>RRID: AB_2315777 |
| Cy3 AffiniPure Donkey Anti-Rabbit IgG (H+L) – 1:200 | Jackson ImmunoResearch Labs | Cat# 711-165-152<br>RRID: AB_2307443 |
| Cy5 AffiniPure Donkey Anti-Guinea Pig IgG (H+L) – 1:200 | Jackson ImmunoResearch Labs | Cat# 706-175-148<br>RRID: AB_2340462 |
| Cy5 AffiniPure Donkey Anti-Mouse IgG (H+L) – 1:400 | Jackson ImmunoResearch Labs | Cat# 715-175-150<br>RRID: AB_2340819 |
| Cy5 AffiniPure Donkey Anti-Rabbit IgG (H+L) – 1:200 | Jackson ImmunoResearch Labs | Cat# 711-175-152<br>RRID: AB_2340607 |

